# Expanding the CRISPR Toolbox with ErCas12a in Zebrafish and Human Cells

**DOI:** 10.1101/650515

**Authors:** Wesley A. Wierson, Brandon W. Simone, Zachary WareJoncas, Carla Mann, Jordan M. Welker, Bibekananda Kar, William A. C. Gendron, Michael A. Barry, Karl J. Clark, Drena L. Dobbs, Maura A. McGrail, Stephen C. Ekker, Jeffrey J. Essner

**Affiliations:** Department of Genetics, Development and Cell Biology, Iowa State University, IA, USA; Department of Biochemistry and Molecular Biology, Mayo Clinic, Rochester, MN, USA

## Abstract

Clustered Regularly Interspaced Short Palindromic Repeats (CRISPR) and CRISPR associated (Cas) effector proteins enable the targeting of DNA double-strand breaks (DSBs) to defined loci based on a variable length RNA guide specific to each effector. The guide RNAs are generally similar in size and form, consisting of a ~20 nucleotide sequence complementary to the DNA target and an RNA secondary structure recognized by the effector. However, the effector proteins vary in Protospacer Adjacent Motif (PAM) requirements, nuclease activities, and DNA binding kinetics. Recently, ErCas12a, a new member of the Cas12a family, was identified in *Eubacterium rectale*. Here, we report the first characterization of ErCas12a activity in zebrafish and human cells. Using a fluorescent reporter system, we show that CRISPR/ErCas12a elicits strand annealing mediated DNA repair more efficiently than CRISPR/Cas9. Further, using our previously reported gene targeting method that utilizes short homology, GeneWeld, we demonstrate the use of CRISPR/ErCas12a to integrate reporter alleles into the genomes of both zebrafish and human cells. Together, this work provides methods for deploying an additional CRISPR/Cas system, thus increasing the flexibility researchers have in applying genome engineering technologies.

## Introduction

CRISPR systems have been widely adopted in zebrafish research due to their efficacy and ease of reprogramming DNA binding activity, which is mediated by a single chimeric short guide RNA (sgRNA) molecule ^1 2 3^. The CRISPR toolbox continues to expand with the identification of systems that display varying PAM requirements and produce a different DSB architecture ^4^. While Cas9 proteins often hydrolyze DNA leaving a blunt-ended cut three base pairs (bps) upstream from the 5’ end of the PAM sequence, Cas12a proteins from *Acidaminococcus* and *Lachnospiraceae* spp. hydrolyze DNA in a staggered fashion, cutting on the 3’ side of the PAM and leaving 5’ single-stranded overhands of 4 nucleotides ^5^. The architecture of the DSB and end resection products are critical determinants of DNA repair pathway activation. ^6^. For example, inducing DNA overhangs with staggered CRISPR/Cas9 nickases targeted to opposite strands can stimulate precision genome engineering using oligonucleotides ^7^. Thus, there is demand for CRISPR variants that generate different DSB architectures and elicit more predictable repair outcomes for precision genome engineering.

CRISPR/Cas12a activity has been reported in zebrafish by injection of sgRNA/Cas12a protein ribonucleoprotein (RNP) complexes ^8, 9^. In these studies it was shown that Cas12a - mediated DNA cleavage could be further enhanced by a 34ºC heat shock or by co-targeting of nuclease dead Cas9 (dCas9) to the Cas12a target site, indicating that DNA melting is a potential rate limiting step for Cas12a. Cas12a resulted in increased efficiency of oligonucleotide incorporation as compared to Cas9 into genomic cut sites by homology-directed repair, suggesting the two enzymes may employ distinct mechanisms and result in different genomic DNA end resection products ^8^. A CRISPR/Cas12a system was identified in *Eubacterium rectale*, and found to be ancestrally related to the type V class II family of CRISPR proteins sharing the greatest similarity with Cas12a from *Acidaminococcus* sp. (AsCas12a) (Figure 1a). *Eubacterium rectale* Cas12a (ErCas12a) is a 1262 amino acid protein that recognizes a 5’-YTTN PAM and uses a 42 or 56 base guide RNA to recognize and catalyze site-specific DNA cleavage^10^. CRISPR/ErCas12a is active in human cells ^10^, but its application for gene targeting *in vivo* and *in vitro* has yet to be described.

**Figure 1.**
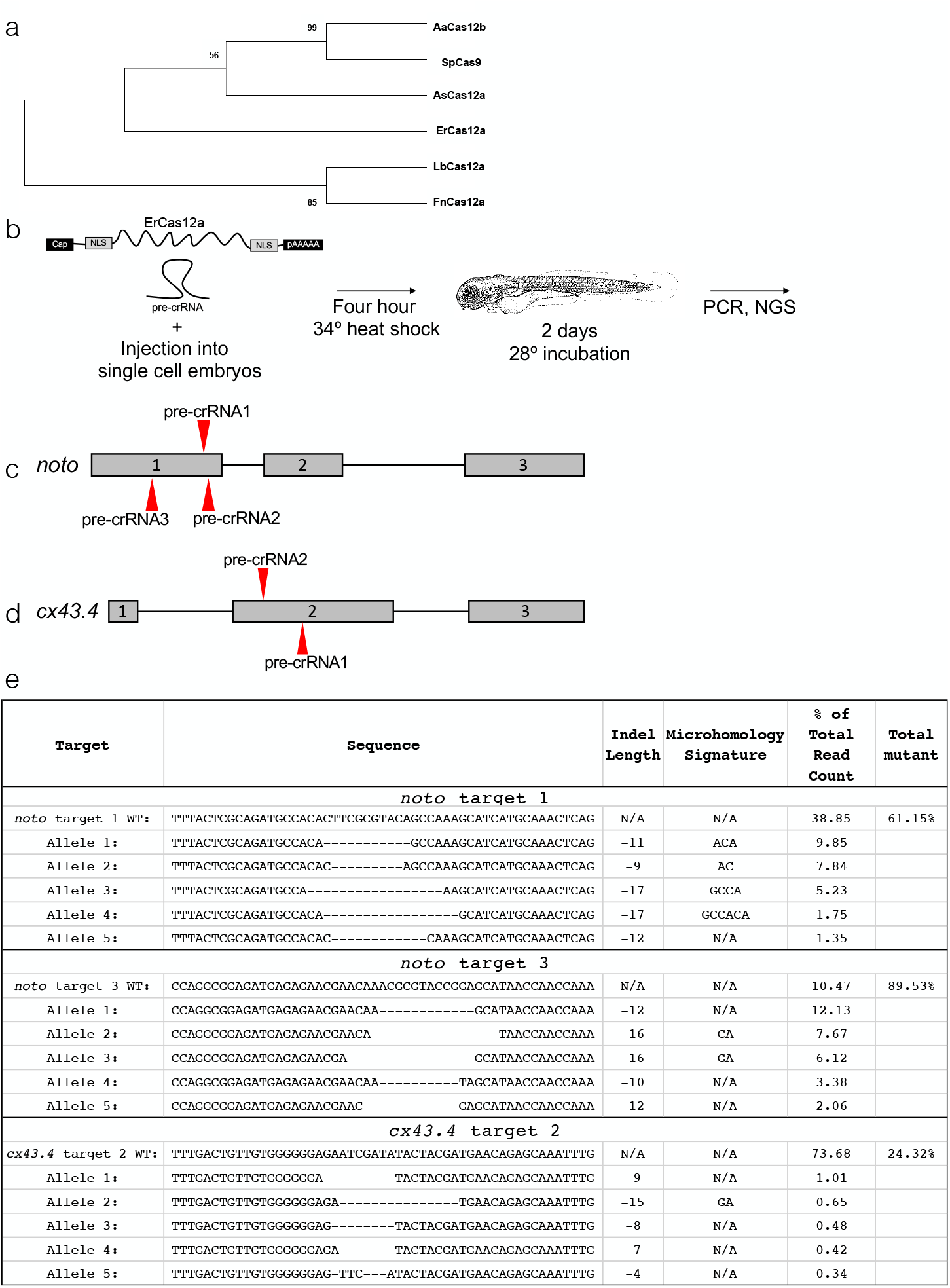
Characterization and activity of CRISPR/ErCas12a in zebrafish. (a) Phylogenetic relationship between known CRISPR associated proteins and ErCas12a. Evaluation of branching was ensured by bootstrap statistical analysis (1000 replications). (b) Workflow showing dual NLS ErCas12a mRNA and pre-crRNA injection into single cell animals. Animals are heat shocked for 4 hours and then allowed to develop normally until DNA is isolated and analyzed. (c) Schematic and sequences of pre-crRNA used to target *noto.* (d) Schematic and sequences of pre-crRNA used to target *cx43.4*. (e) Results displaying the wildtype and top 5 mutated alleles after AmpliconEZ analysis of two *noto* pre-crRNAs and one *cx43.4* pre-crRNA.

Microhomology mediated end joining (MMEJ) is a DNA repair pathway that uses short regions of homology between resected DNA ends to drive repair, leading to predictable outcomes after nuclease targeting ^11 12 13^. MMEJ-based gene editing and targeted integration using MMEJ has been described in zebrafish and mammalian cells, although the length of the homology arms appear to be more similar in length to those employed in single strand annealing (SSA) ^14 15 16 17^. To use MMEJ/SSA for gene targeting, nuclease targeting of a donor plasmid *in vivo* liberates a donor cassette, exposing homology arms that are complementary to the nuclease cut site in the genome, and thereby directs integration into genomic DNA. A more general term for using this strategy for gene targeting is Homology-Mediated End Joining (HMEJ) ^18^. We recently reported high efficiency gene targeting by HMEJ with short homology arms called GeneWeld ^19^. GeneWeld involves the simultaneous cleavage of a donor plasmid and a genomic target with designer nucleases to reveal 24 or 48 bp homology arms that can be used by the cellular MMEJ or SSA machinery for homology directed repair leading to donor integration. GeneWeld works with genomic DSBs generated by either TALENs or CRISPR/Cas9, but other nucleases have not yet been tested.

Here, we report the successful application of ErCas12a for gene targeting in zebrafish, widely used to model development and disease, and in human cells. Consistent with its similarity to other Cas12a proteins, high ErCas12a activity requires a 34ºC heat shock treatment in zebrafish. However, injection of mRNA encoding for ErCas12a is sufficient to induce DSB activity, and pre-crRNA can serve as an effective RNA guide for ErCas12a, in contrast to reports using previously described Cas12a and related systems ^8, 20^. We developed a Universal pre-crRNA (U-pre-crRNA) for ErCas12a which can be used in both zebrafish and human cells for donor DNA cleavage and show that CRISPR/ErCas12a potently induces strand annealing mediated repair (SAMR) in a genomic reporter locus at efficiencies greater than CRISPR/Cas9. Additionally, CRISPR/ErCas12a promotes GeneWeld activity at rates similar to CRISPR/Cas9. Finally, we apply GeneWeld with ErCas12a in human cells, and demonstrate ErCas12a mediated targeted integration activity at several human loci, including the safe harbor locus *AAVS1*.

## Results

### ErCas12a induces indels in zebrafish embryos

We used informatic analyses to categorize a new potential gene editor (GenBank Accession MH347339.1) that aligns as a novel Cas12a protein, ErCas12a (Figure 1a, Supplemental Figure 1). To assess the activity of ErCas12a, we first synthesized a codon optimized version for expression in eukaryotes and added dual SV40 nuclear localization signals at the N- and C-termini, as dual NLS was shown to increase efficacy of Cas9 in zebrafish ^4^. To test ErCas12a gene editing activity *in vivo*, ErCas12a mRNA was co-injected with a pre-crRNA into single cell zebrafish embryos followed immediately by heat shock at 34°C for 4 hours (Figure 1b). Three pre-crRNAs and one crRNA were designed to target exon 1 of the zebrafish *noto* gene (Figure 1c). Co-injection of ErCas12a mRNA with *noto*-pre-crRNA1 and incubation at 28ºC did not result in gene editing activity, similar to previous reports using Cas12a (Moreno-Mateos et al., 2017^8^ and data not shown). However, injection of ErCas12a mRNA with *noto*-pre-crRNA1 or *noto*-pre-crRNA3 followed by a 4-hour 34ºC heat shock resulted in phenotypes characteristic of biallelic loss of *noto*, including loss of the notochord and a shortened tail ^21^, at efficiencies ranging from 7-61% (Supplemental Figure 2a). PCR across these individual target sites and subsequent heteroduplex mobility shift assays revealed the presence of indels characteristic of repair by NHEJ activity (Supplemental Figure 2b). *noto*-pre- crRNA2 and *noto-*crRNA1 did not elicit a phenotype, and heteroduplex mobility shift assays across those targets indicate they are inactive sgRNAs (data not shown). Efficiencies of biallelic *noto* inactivation at the same target varied considerably from injection to injection (Supplemental Figure 2), consistent with previous reports in human cells and zebrafish using CRISPR/Cas12a^22 23 8 20^. To confirm that our results were not specific to a single locus, we injected ErCas12a mRNA and pre-crRNA to target two sites in the *cx43.4* gene (Figure 1d). PCR across the individual target sites and subsequent RFLP analysis demonstrated that *cx43.4*-pre-crRNA2 was active (Supplemental Figure 2d), while *cx43.4-*pre-crRNA1 was inactive (data not shown).

To gain a better understanding of the frequency of ErCas12a insertion/deletion (indel) events, we used Illumina next generation sequencing (NGS) to analyze the efficiencies of mutation at the active *noto* and *cx43.4* target sites. DNA amplicons from 5 *noto* mutant embryos injected with ErCas12a mRNA and *noto-*pre-crRNA1 or *noto-*pre-crRNA3 and 5 *cx43.4* pre-crRNA1 injected embryos were selected randomly for next generation sequencing. As expected for biallelic inactivation at *noto*, ~61% of alleles at *noto*-pre-crRNA1 showed indels, characteristic of NHEJ and/or MMEJ after nuclease targeting (Figure 1e). The more efficient *noto-*pre-crRNA3 showed ~90% of alleles containing indels (Figure 1e). However, in agreement with the RFLP analysis, the majority (~74%) of alleles sequenced at *cx43.4* had the wild type sequence, indicating that not all targeting events with ErCas12a are efficient enough to produce biallelic mutation of the target locus (Figure 1e). Taken together, these data indicate CRISPR/ErCas12a introduced as an RNA system can be active in zebrafish with a 34ºC heat shock, and that pre-crRNA can serve as an active RNA guide for directing ErCas12a activity to genomic target sites.

### ErCas12a elicits strand annealing mediated DNA repair more efficiently than SpCas9 in a genomic reporter system in zebrafish

We next employed a stably integrated red fluorescent protein (RFP) reporter system to visually compare the efficiencies with which CRISPR/Cas9 and CRISPR/ErCas12a differentially elicit DNA repair using strand-annealing mediated repair (SAMR). We used GeneWeld to create a transgenic line of zebrafish that contain a single copy of an SAMR reporter, *noto:*RFP-DR48 (Figure 2a; Supplemental Figure 3). The RFP coding sequence is disrupted by a target cassette engineered with Cas9 and ErCas12a sites that is flanked by 48 bp of direct repeat sequence. A double strand break and end resection in the target liberates the 48 nucleotide direct repeats which can restore the RFP reading frame after SAMR, leading to a semi-quantitative read-out of repair efficiencies. Injection of single-cell *noto:*RFP-DR48 transgenic embryos with Cas9 mRNA and UgRNA results in RFP expression in the notochord, indicative of SAMR after CRISPR/Cas9 targeting (Figure 2b).

**Figure 2.**
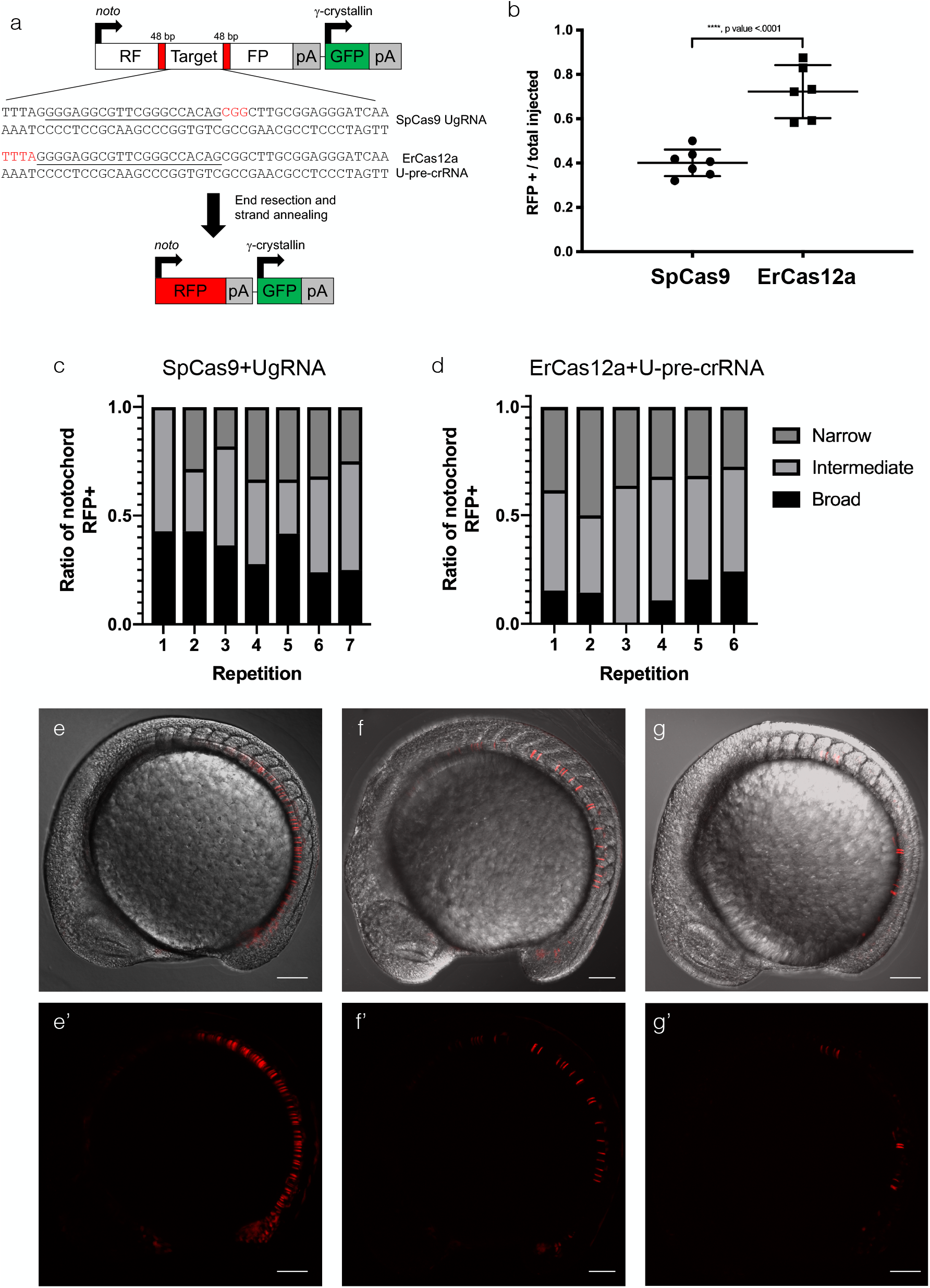
Using *noto:*RFP-DR48 to assay the propensity of ErCas12a and SpCas9 to elicit strand annealing in zebrafish. (a) Schematic of *noto*:RFP-DR48 showing the location of the 48 bp direct repeats flanking both the SpCas9 and ErCas12a universal RNA cursor sites (underline) and PAM sites (red text). (b) Data plot showing the ratio of injected animals displaying RFP in the notochord out of total carrying the transgene. Data plot represents represent mean +/− s.d. p values calculated with one-tailed Student’s t-test. (c) Qualitatively scored ratios of notochord converted to RFP+ after SpCas9 induced SAMR. (d) Qualitatively scored ratios of notochord converted to RFP+ after ErCas12a induced SAMR. (e-g) Representative embryos for broad, intermediate, and narrow conversion of RFP in the notochord.

We then tested whether CRISPR/ErCas12a promotes SAMR in the *noto:*RFP-DR48 assay. We designed a universal pre-crRNA (U-pre-crRNA), with no predicted off target sites in zebrafish or human cells, to direct ErCas12a activity to *noto:*RFP-DR48. Injection of ErCas12a mRNA and U-pre-crRNA resulted in RFP expression in the notochord (Figure 2b). Based on the percentage of injected animals with RFP+ notochords, ErCas12a elicited SAMR at a higher frequency than SpCas9 (Figure 2b). Injected embryos were highly mosaic and qualitatively sorted into 3 classes of notochord RFP expression pattern: broad, intermediate, and narrow (Figure 2e-i). While ~70% of ErCas12a-injected animals showed RFP+ cells in the notochord as opposed to ~40% in SpCas9-injected embryos, ErCas12a repair events are equally mosaic (Figure 2f). These results indicate that *noto:*RFP-DR48 is a viable assay for screening the propensity of designer nucleases to elicit SAMR and that the enzymatic activity of ErCas12a enhances activation of SAMR over Cas9 *in vivo*.

### Using ErCas12a for precise integrations in zebrafish

We leveraged the activity of the U-pre-crRNA to determine whether CRISPR/ErCas12a can catalyze targeted integration of fluorescent reporters using GeneWeld ^19^. First, we injected nErCas12an, *noto*-pre-crRNA1, U-pre-crRNA, and a GFP reporter plasmid programmed with the U-pre-crRNA target site and 24 bp of homology 5’ to the noto-crRNA1 genomic site to promote gene targeting at *noto* (Supplemental Figure 4a, Supplemental Table 4). Fluorescent positive notochord cells were observed at the 18-somite stage, indicating in frame integration of the GFP cassette at *noto* (Supplemental Figure 4b, b’). On average, 24% of embryos injected display targeted *noto* integrations (Supplemental Figure 4c). However, notochords were highly mosaic, indicating a low efficiency of somatic integration activity in tissue types that form the notochord (Supplemental Figure 4b, b’). GFP was often observed outside of the notochord, yet in the mesoderm, as expected with biallelic disruption of *noto* and transfating of notochord cells ^21^. Junction fragment PCR was conducted on GFP+ embryos to confirm that integration was directed by the programmed homology. The expected PCR band was recovered in GFP+ embryos, while no PCR band was detected in the control experiment performed without the genomic pre-crRNA (Supplemental Figure 4d, 4e). DNA sequencing confirmed precise junctions at the 5’ end of the integration (Supplemental Figure 4f).

We next designed a GeneWeld donor with both 5’ and 3’ homology domains flanking the GFP cassette to promote precise repair at both sides of the integration site (Figure 3a) ^19^. Targeting *noto* using pre-crRNA3 with GeneWeld resulted in an average of 31% of embryos with GFP+ notochords (Figure 3b, 3b’, 3c; Supplemental Table 4). Most notochords were highly mosaic and displayed GFP expression indicative of lost cell fate, as in *noto* target 1 targeting (Supplemental Figure 4b), although some events were recovered in which >90% of the notochord expressed GFP (Figure 3b). Predicted 5’ and 3’ junction fragments were recovered by PCR in GFP+ embryos (Figure 3d, 3e), and junction fragment sequencing from a single embryo demonstrated precise integration at both ends of the cassette (Figure 3f). These data indicate that ErCas12a can be an effective nuclease for catalyzing integration in zebrafish using GeneWeld, but that optimization is needed to enhance somatic integration efficiency to levels comparable to those obtained using Cas9.

**Figure 3.**
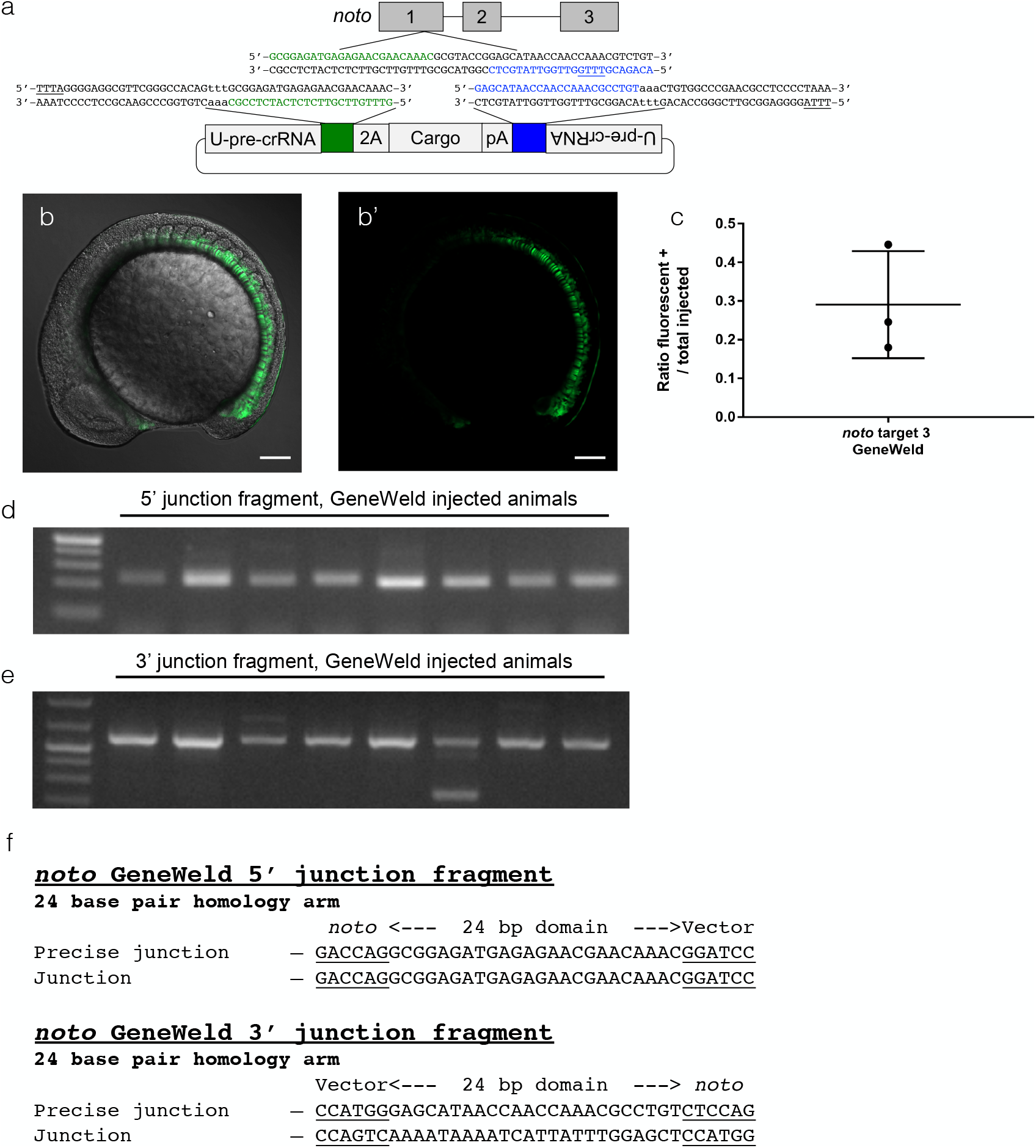
Targeting *noto* with GeneWeld. (a) Schematic of *noto* showing designed homology for precise integration using ErCas12a and the U-pre-crRNA. Green is designed 5’ homology. Blue is designed 3’ homology. The PAM for ErCas12a targeting in the genome and donor is underlined. (b, b’) Confocal Z-stack image showing broad GFP expression in the embryo. Scale bar is 100 um. (c) Data plot showing the ratio of embryos with GFP expression in the notochord out of total injected embryos. Data plot represents mean +/− s.d. (d) Gel showing 5’ junction fragment expected after precise integration using GeneWeld. (e) Gel showing 3’ junction fragment expected after precise integration using GeneWeld. (f) DNA sequencing of lane 5 in (d) and (e) showing a precise integration using the programmed homology.

### ErCas12a induces double-strand breaks in human cells

We developed an all-in-one vector for expression of the dual nuclear localization signal ErCas12a and a pre-crRNA *in vitro* (Figure 4a). We targeted two therapeutically relevant “safe harbor” loci, *AAVS1* and *CCR5* ^24^ (Figure 4b, d). We additionally targeted the *TRAC* locus due to its reported significance in generating Chimeric Antigen Receptor T cells (Figure 4e) ^25^. *in vitro* cleavage activity of ErCas12a with our expression system was assessed using a T7 endonuclease (T7EI) assay to determine whether a DSB and subsequent indels were induced at the *AAVS1* locus after transfection into HEK293 cells. The expected banding pattern was apparent in DNA amplified from pErCas12a-AAVS1-pre-crRNA1 transfected cells but not control cell DNA, indicating that ErCas12a was active at the AAVS1 target site (Figure 4c). No indel activity was detected at the *AAVS1* pre-crRNA2 site by T7E1 assay (data not shown). These data suggest that the all-in-one expression system is functional *in vitro*.

**Figure 4.**
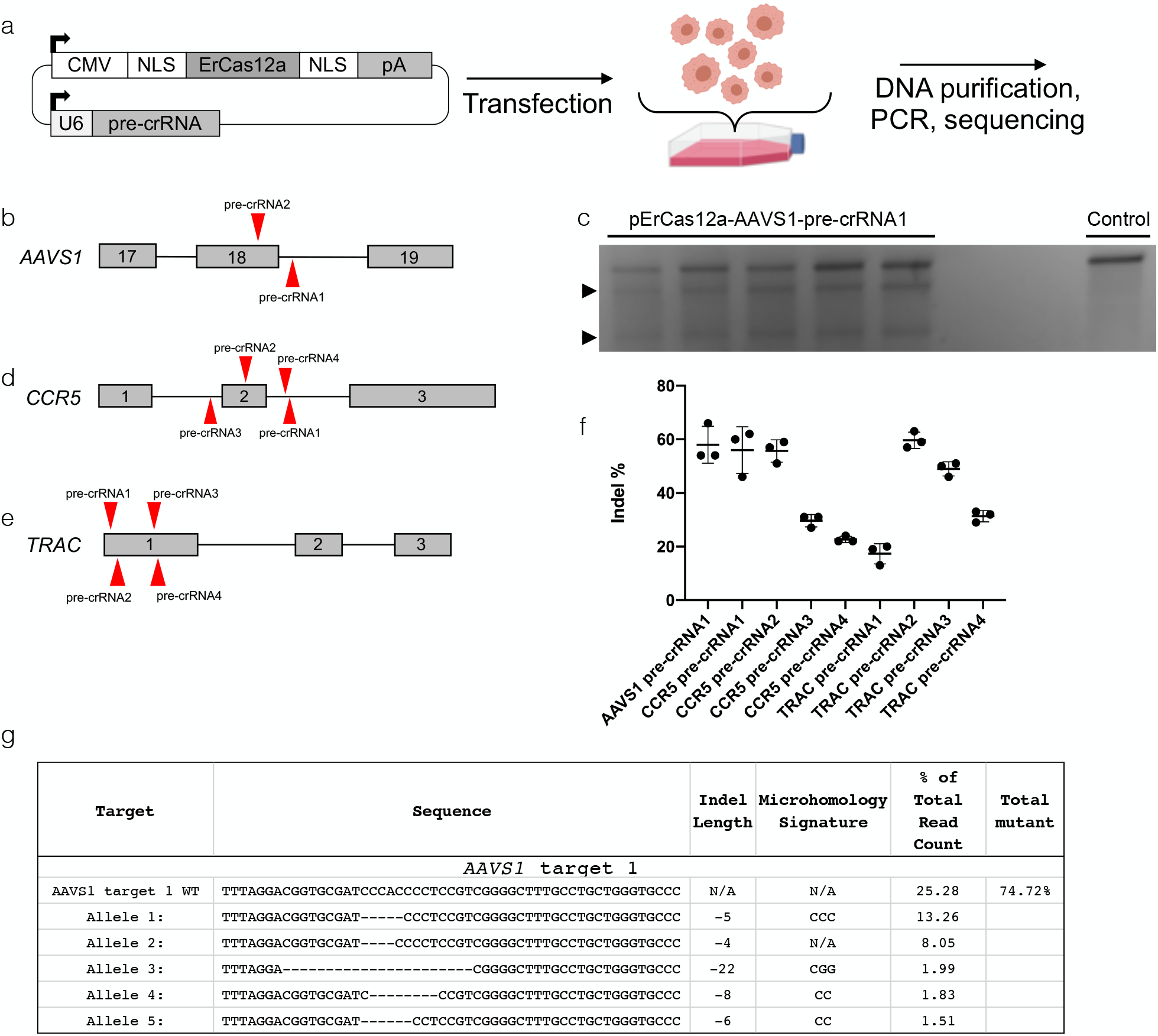
ErCas12a activity in human HEK293 cells. (a) Schematic showing the workflow for ErCas12a and pre-crRNA expression in human cells from a cis expression plasmid. After transfection, cells are recovered and then analyzed. (b) Schematic of two pre-crRNAs designed to target *AAVS1*. (c) T7E1 assay showing *AAVS1-*pre-crRNA1 is active. Arrowheads are cleaved bands, indicating nuclease activity. (d) Schematic of four pre-crRNAs designed to target *CCR5*. (e) Schematic of 4 pre-crRNAs designed to target *TRAC*. (f) Graph showing percentage of indels after using pre-crRNAs to target *AAVS1, CCR5*, and *TRAC* in HEK293 cells as determined by ICE analysis. Data plot represents mean +/− s.d. (f) Results displaying the wildtype and top 5 mutated alleles after AmpliconEZ analysis of *AAVS1* targeted DNA.

To gain a quantitative understanding of the cleavage activity of ErCas12a at the *AAVS1, CCR5, TRAC* target sites, PCR amplicons were submitted for Sanger sequencing and analyzed using ICE software, which infers CRISPR activity from sequencing trace reads ^26^. ICE analysis showed targeting *AAVS1* with ErCas12a, and pre-crRNA1 created indels in 58% of sequenced amplicons (Figure 4f, Supplementary Table 6). Targeting *CCR5* with ErCas12a at four sites resulted in a range of mutagenesis efficiency from 20-60% (Figure 4f, Supplementary Table 6). Similarly, we tested 4 pre-crRNAs targeting *TRAC* and found the most efficient resulted in 60% of sequenced amplicons contained indels (Figure 4f, Supplementary Table 6). Because ICE is based on Sanger sequencing data, which considers fewer reads than NGS, it is likely to underestimate the true percentage of indels. We therefore measured indel activity at the *AAVS1*-pre-crRNA1 target site using Illumina sequencing, which showed that ~74% of recovered alleles were edited (Figure 4g). Taken together, these data demonstrate that ErCas12a is active in human cells at clinically relevant genomic loci.

### ErCas12a elicits strand annealing mediated DNA repair more efficiently than SpCas9 in an episomal reporter system in human cells

Episomal reporters have been used routinely in cell culture systems to assay DNA repair pathways and identify the proteins involved ^27 28 29 30^. We modified the *noto*:RFP-DR48 cassette for use as a reporter in mammalian cell culture, creating pMiniCAAGs:RFP-DR48 (Figure 5a). As observed in zebrafish, transfection of pMiniCAAGs:RFP-DR48 with a nuclease and requisite sgRNA into human cells is expected promote a double-strand DNA break at the nuclease target site, resulting in a fluorescent readout of nuclease activity if SAMR is used to repair the double strand break. Transfection of HEK293 cells with pMiniCAAGs:RFP-DR48 and pErCas12a-U-pre-crRNA or pCas9-UgRNA resulted in RFP+ cells, demonstrating efficient SAMR with both ErCas12a and Cas9 nucleases (Figure 5a). Quantification of RFP+ cells by flow cytometry indicated SAMR induction was significantly greater (*p* < 0.0044) with ErCas12a compared to SpCas9 (Figure 5b, Supplemental Figure 5). Together, these results show ErCas12a nuclease has similar cleavage activity in human cells and zebrafish, and is more efficient than Cas9 in mediating SAMR in *vivo* and *in vitro*.

**Figure 5.**
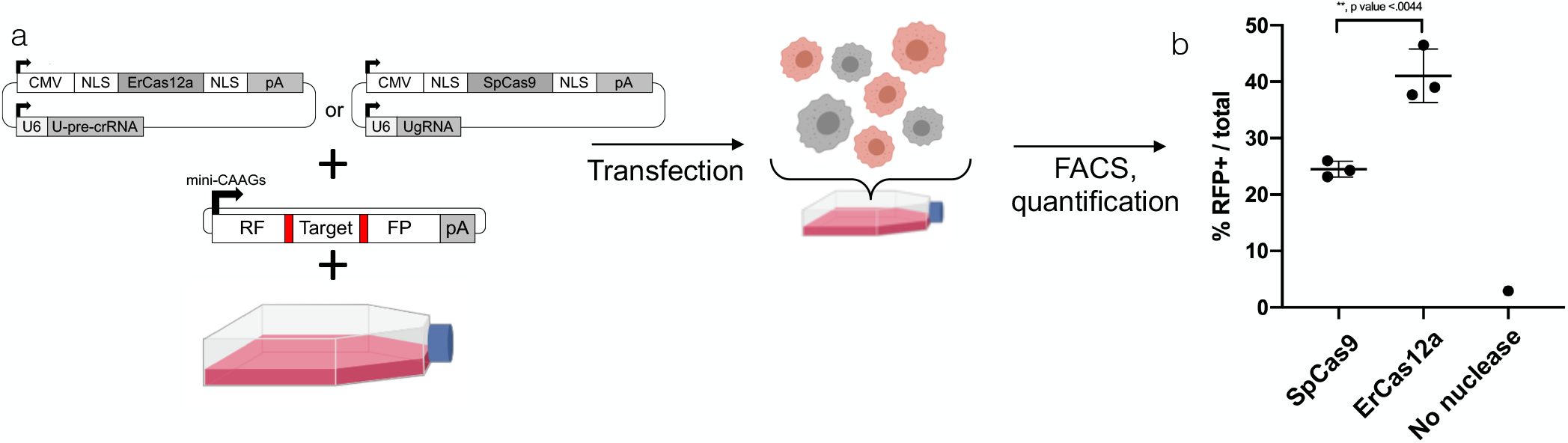
Using pMini-CAAGs::RFP-DR48 to assay the propensity of ErCas12a and SpCas9 to elicit strand annealing in HEK293 cells. (a) Schematic showing the workflow for determining strand annealing mediated repair in human cells. An all in one expression plasmid for ErCas12a or SpCas9 is transfected along with pMini-CAAGs::RFP-DR48. Cells are allowed to recover and then FACS sorted for RFP+ cells. (b) Quantification of RFP+ cells per total cells analyzed by flow cytometry. Data plots represent mean +/− s.d. p values calculated with one-tailed Student’s t-test.

### ErCas12a promotes precise integration in human cells

Because of *AAVS1*’s utility in therapeutic expression of transgenes, we next wanted to determine whether ErCas12a could mediate precise transgene insertion at the pre-crRNA1 site in *AAVS1* ^31 32 33^. To test this, we used the GeneWeld strategy to mediate integration of a pCMV:GFP::Zeo reporter containing 48 bp 5’ and 3’ arms homologous to the *AAVS1* pre-crRNA1 site (Figure 6a, b). HEK293 cells were transfected with the pCMV:GFP:Zeo donor plasmid and the all-in-one pErCas12a-AAVS1-U plasmid that expresses ErCas12a, *AAVS1* pre-crRNA1, and U-pre-crRNA (pErCas12a-AAVS1-U). Transfections were also done with the pCMV:GFP:Zeo-48 donor plasmid and pErCas12a-AAVS1 pre-crRNA1 without the U-pre-crRNA, or pCMV:GFP:Zeo-48 donor plasmid alone. GFP+ cells were isolated by fluorescence activated cell sorting (FACS) one day post electroporation to control for plasmid delivery, and then screened for stable integration by flow cytometry 2 weeks post transfection to measure dropout of GFP+ cells (Figure 6c, Supplemental Figure 6). The data revealed a significant 2.5 fold increase (p<0.0003) in the number of cells that remained GFP+ when transfected with donor molecule, ErCas12a and the pre-crRNA, as compared to donor alone control (Figure 6d, Supplemental Figure 6b-6c’’). GFP+ cell populations were screened for targeted integration at the *AAVS1* pre-crRNA1 site by junction fragment PCR and sequencing. The 5’ junction analysis showed precise repair at the integration site in the cell population transfected with pErCas12a-AAVS1-U and the donor (Figure 6e, f) but was absent in the donor alone population. These data suggest ErCas12a is a tractable nuclease for GeneWeld-mediated precision integration at the *AAVS1* safe harbor locus in human cells.

**Figure 6.**
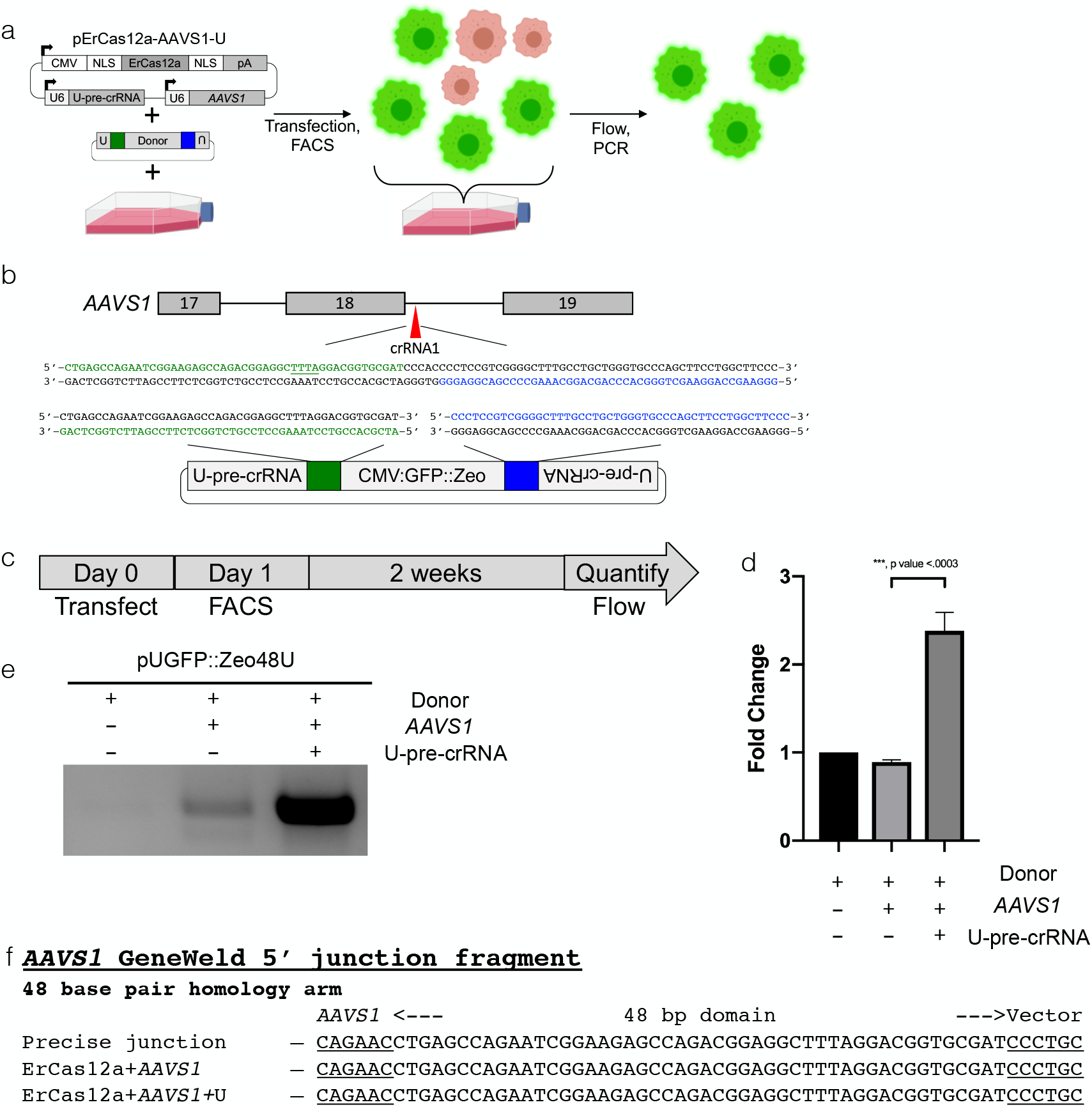
Using ErCas12a for targeted integration in human cells. (a) Schematic showing the workflow for targeted integration at *AAVS1* in HEK293 cells. (b) Schematic of *AAVS1* showing designed homology for precise integration using ErCas12a and the U-pre-crRNA to liberate the cassette. Green block, 5’ homology; Blue block, 3’ homology. The PAM for ErCas12a targeting in the genome and donor is underlined. (c) Schematic for flow analysis after transfection of GeneWeld reagents for targeting *AAVS1*. (d) Bar graph showing the ratio of cells with GFP expression as determined by flow cytometry. Graph represents mean +/− s.d. p values calculated with one-tailed Student’s t-test. (e) Junction fragment gel showing the expected 5’ integration amplicon. No amplicon is seen when transfecting donor alone. (f) DNA sequencing of lanes in (e) showing a precise integration using the programmed homology.

## Discussion

In this report, we establish that CRISPR/ErCas12a is an active nuclease in both zebrafish and human cells and can serve as an efficient alternative to CRISPR/Cas9 for generating HMEJ-mediated gene targeting events *in vivo*. Targeting in zebrafish embryos, we created frame shift indel alleles in the *noto* and *cx43.4* at up to ~90% and ~24%, respectively. ErCas12a activity is temperature-dependent in zebrafish and requires a 34ºC heat shock for activity. In a genomic reporter assay, ErCas12a elicits SAMR in a reporter assay at levels nearly 2-fold greater than Cas9 in both zebrafish and human cells. ErCas12a is also a viable nuclease for mediating gene targeting using the GeneWeld strategy in zebrafish and human cells. We detected robust double-strand break induction at three therapeutically relevant human loci and demonstrated ErCas12a is effective for directing GeneWeld integrations at the *AAVS1* locus in human cells. These results suggest that ErCas12a could be employed as a tractable nuclease for gene editing across multiple species for both basic research and clinical applications.

In prior reports of CRISPR/Cas12a activity in zebrafish, appreciable nuclease activity was only observed after RNP delivery and heat shock, “proxy-CRISPR” to relax chromatin structure, or targeting a gene with multiple crRNAs simultaneously ^8 20^. Cas12a activity was shown to be dependent on the stability of the bound pre-crRNA or crRNA and although pre-crRNA was ineffective for Cas12a nuclease activity under standard conditions, longer heat shocks rescued activity, indicating a likely kinetics issue with the use of pre-crRNA. In our experiments, pre-crRNA injection with ErCas12a mRNA permitted nuclease activity, highlighting a difference between other Cas12a related proteins and ErCas12a, either in their ability to stably bind the RNA guide and access DNA, or in the stability of their respective pre-crRNA structures. Because the use of multiple crRNAs per target gene in tandem dramatically increases the possibility of off target effects, it is notable that in this study, ErCas12a displays efficient nuclease activity with only a single pre-crRNA per target gene.

While ErCas12a showed enhanced activation of SAMR in comparison with Cas9 in our *noto*:RFP-DR48 assay, targeted integration via GeneWeld using ErCas12a in zebrafish displayed no substantive difference compared to our previous report using Cas9 ^19^. The increases SAMR frequency detected using the reporter assay may be because repair of DNA *in cis* is more efficient than *in trans*, and thus the outcome of strand annealing in a genomic reporter will not translate to targeted integrations.

Reported levels of mosaicism in gene targeting using HMEJ in zebrafish vary greatly in the literature ^15 19^. Although ErCas12a is an effective catalyst for GeneWeld integrations, mosaicism of expression is qualitatively higher than when using Cas9 as the GeneWeld nuclease ^19^. Consistent with the observation that SAMR in *noto*:RFP-DR48 is mosaic, it was reported recently that translation of Cas9 mRNA and subsequent gene editing after one-cell stage injection is not complete until the 16 or 32 cell stage, while RNP injections result in appreciable nuclease activity by the 2-4 cell stage ^34^. Thus, *noto*:RFP-DR48 is an assay system where injection conditions can be optimized to enhance somatic gene targeting and decrease mosaicism in cell types that arise from the mesoderm. Also, *noto*:RFP-DR48 represents a novel reporter in zebrafish for determining strategies for altering DSB repair pathway choice, such as using small molecules or dominant-negative proteins.

Previously reported Cas12a orthologs have displayed relatively poor editing efficiency in mammalian systems ^35 22 36 23^. AsCas12a and LbCas12a are among the best characterized Cas12a systems but have very modest activity in human cells ^22, 36^. To address this issue, additional Cas12a systems with enhanced activity and differential targeting ability such as FnCas12a were engineered and characterized. Although FnCas12a uses a more common TTN PAM sequence, its cleavage activity is still modest ^35^. Editing efficiency of the Cas12a systems has since been enhanced by engineering the crRNA to increase activity without a loss of specificity ^37^. Additionally, recent work has identified particular point mutations in Cas12a variants that enhance the targeting range, activity, and fidelity of the nuclease ^38^. This suggests the possibility of a future ErCas12a variant in which cleavage kinetics could be leveraged with broadened PAM targeting to target regions typically restricted to SpCas9. Going forward, it will be interesting to evaluate the effects of various modifications of crRNAs and the protein itself to optimize mutagenesis and gene targeting.

The 5 base thymidine stretch in the secondary structure of the ErCas12a crRNA could cause cryptic termination of guide expression ^39^ and lead to decreased expression of the guide RNA, potentially decreasing nuclease activity. Testing crRNA designs that remove this thymidine stretch may increase crRNA expression and nuclease efficiency. It has been shown previously that a single U6 promoter is sufficient to express multiple crRNAs with Cas12a for multiplex gene editing due to the intrinsic self-processing activity of the nuclease ^2^. It is of interest for future investigations to determine if ErCas12a has similar properties in order to leverage it for multiplex gene editing.

Interestingly, the RuvC domain in Cas12a family members is sufficient to elicit indiscriminate ssDNA and ssRNA cleavage ^40^. Though ErCas12a shares only 31% sequence homology with Cas12a orthologue AsCas12a (Supplemental Figure 1) and the RuvC domain is conserved among Cas12a orthologues and ErCas12a, it is possible that long range interactions at the catalytic pocket differ in ErCas12a due to the relatively low amino acid similarity. Therefore, further studies are needed to elucidate if ErCas12a shares similar nonspecific nuclease activity.

Though Cas12a systems has been shown to be amendable to HDR-mediated integration, these transgene integration events have been shown to be highly dependent on using RNP complexes in concert with modified crRNAs to show appreciable activity ^23^. Here, we demonstrate ErCas12a’s utility in generating robust editing efficiency without chemically modified crRNAs, showing its utility in streamlined and accessible gene editing for most research applications. Further, we show ErCas12a mediates precise integration without engineering of the nuclease or the corresponding guide RNA with mRNA and plasmid DNA.

While we show that ErCas12a is compatible with GeneWeld. We also demonstrated that in the absence of the UgRNA liberating the donor there is still integration of the transgene at *AAVS1.* When GeneWeld is used with Cas9, the donor homology arms can include a portion of the genomic target but exclude the 3’ PAM sequence and prevent nuclease targeting at the homology arms. However, because ErCas12a utilizes a 5’ PAM and induces a distal DSB, it is likely that the homology arm is being targeted by ErCas12a and linearizing the donor at the 5’ end. In our design the homology arms contain up to 17bp of target region that may be sufficient for a Cas12a-like seed region for the crRNA and facilitate a DSB ^40 41^. With further optimization of both the crRNA and donor constructs, ErCas12a can readily be adapted for precise integration in mammalian systems.

## Conclusion

Alternative nuclease systems beyond canonical CRISPR/Cas9 are of interest in the precision therapeutics field as they expand the available toolbox for precision gene editing. Here, we demonstrate effective genome editing in both zebrafish and human cells using the newly described CRISPR/ErCas12a system, which increases the number of accessible regions in the genome due to its AT-rich PAM sequence. In zebrafish, CRISPR/ErCas12a promotes efficient somatic mutation and HMEJ-mediated precise integration of donor cassettes. In human cells, CRISPR/ErCas12a efficiently targeted therapeutically relevant loci, including the safe harbor loci *AAVS1* and *CCR5*, and facilitated transgene integration. Our data suggest that, with further optimization, ErCas12a may be an invaluable tool for gene editing in both basic research and future clinical gene therapy applications.

## Author Contributions

WAW, BWS, SCE, JJE conceived the study. WAW designed nlsErCas12anls and conducted the zebrafish work. BWS conducted the human cell work. The manuscript was written by WAW and BWS with input and edits from ZWJ, DD, KJC, SCE, MM and JJE. JMW created the vector backbone for *noto*:RFP-DR48. CM analyzed the NGS data. WAG and MAB designed the *in vitro* knock-in backbone. BK conducted the Cas protein phylogenetic analysis. All authors approved of the manuscript and signed off on the study.

## Acknowledgements

The authors would like to acknowledge Inscripta for providing their *Eubacterium rectale* ErCas12a (Mad7) sequence as an open source nuclease for the gene editing community. pX601-AAV-CMV::NLS-SaCas9-NLS-3xHA-bGHpA;U6::BsaI-sgRNA is a gift from Dr. Feng Zhang. Dr. Hirotaka Ata for experimental design discussions, Dr. Karthik Murugan for providing insight into *in vitro* knock in experimental results, and Gabriel Martinez Galvez in assisting in analyzing NGS data. We would further like to thank the Mayo Clinic Microscopy and Cell Analysis Core for conducting flow cytometry and fluorescence activated cell sorting. This work was supported by NIH grants OD020166, GM63904 and the Mayo Foundation.

## Materials and methods

### Zebrafish husbandry

Zebrafish were maintained in Aquatic Habitats (Pentair) housing on a 14 hour light/10 hour dark cycle. Wild-type WIK were obtained from the Zebrafish International Resource Center. All experiments were carried out under approved protocols from Iowa State University IACUC.

### nErCas12an cDNA cloning

gBlocks were ordered from IDT with zebrafish codon optimized ErCas12A cDNA sequences based on Inscripta public disclosure and the addition of dual NLS sequences at the 5’ and 3’ end of the cDNA (Supplemental Table 1). Three gBlock dsDNA templates were amplified with KOD HotStart DNA polymerase (EMD Millipore) using primers ErCas12af1/ErCas12aAr1, ErCas12af2/ErCas12ar2, ErCas12af3/ErCas12ar3, cut with respective restriction enzymes, and four-part restriction cloning into NcoI/SacII cut pT3TS vector backbone was performed (Plasmid #46757 Addgene). Sequence of ERCAS12A using primer walking and a full annotation was confirmed.

### Injection protocol

Linear, purified pT3TS-nCas9n or pT3TS-nErCas12an was used as template for in vitro transcription of capped, polyadenylated mRNA with the Ambion T3TS mMessage mMachine Kit. mRNA was purified using Qiagen miRNeasy Kit. The Cas9 universal sgRNAs were generated using cloning free sgRNA synthesis as described in Varshney et al., 2015 and purified using Qiagen miRNeasy Kit. All ERCAS12A pre-crRNA and crRNA was ordered as custom RNA oligos from Synthego with sequences described in Figure 1a.

### Heat shock protocol

Immediately after injection, embryos were placed in a 34ºC incubator for 4 hours. At 4 hours, embryos were sorted for fertilization and fertilized embryos were moved to 28ºC incubator as normal.

### DNA isolation and PCR genotyping

Genomic DNA for PCR was extracted by digestion of single embryos in 50mM NaOH at 95ºC for 30 minutes and neutralized by addition of 1/10^th^ volume 1M Tris-HCl pH 8.0. GoTaq Green was used as DNA polymerase master mix with the primers listed in Supplemental Table 2. AmpliconEZ from GeneWiz was used for NGS sequencing (see below) using primers ErCas12anoto1fEZ and ErCas12anoto1rEZ for *noto* target 1, ErCas12anoto3fEZ and ErCas12anoto3rEZ for *noto* target 3, and cxm7gRNA2fEZ, cxm7gRNA2rEZ for *cx43.4* listed in Supplemental Table 2. GFP+ embryo 5’ junction fragments for *noto* target 1 and target 3 were PCR-amplified with primer notojxnf and gfp5’r listed in Supplemental Table 2. GFP+ embryo 3’ junction fragments for *noto* target 3 were PCR-amplified with primer GFP3’F and notojxnr listed in Supplemental Table 2. All junction fragment products were cloned into pCR4-TOPO vector and sequenced (Invitrogen).

### Donor vector preparation

Donor vectors were prepared and purified as described previously (Wierson et al., 2018). Homology arms are built as follows: One arm begins 13 bp 3’ of the PAM while the other arm begins immediately outside of the 3’ end of the crRNA target site. See Figure 3 and Figure 4 for homology arm design, and Supplemental Table 3 for GeneWeld homology arm oligos used for Golden Gate cloning. Gene targeting oligos and donor vector sequences are listed in Supplemental Tables 2 and 3.

### Generating *noto:*RFP-DR48

Zebrafish RFP assay generation, injection, and line isolation

pPRISM-V3 was PCR amplified with v3f and v3r to remove the ocean-POUT terminator and add SgrAI and SpeI cloning sites (Supplemental Table 2). Bactin 3’ UTR was PCR amplified using KOD polymerase with primers bactinf and bactinr to add SgrAI and SpeI enzyme sites for sticky end cloning. pPRISM-V3(pout negative) amplicon and bactin 3’ UTR were cut with SgrAI and SpeI. After ligation with Fisher Optizyme T4 Ligase and sequence verification to create pPRISM-V3-bactin, pPRISM-V3-bactin-SSA-DR48 was created by simultaneously adding the NBM and creating a direct repeat with phosphorylated primers DRf and DRr using pPRISM-V3-bactin with KOD polymerase followed by Fisher Optizyme T4 Ligation and sequence verification. To target these constructs to *noto*, homology domains up and downstream of a genomic CRISPR/Cas9 target site were chosen as described in Wierson et al., 2018. Oligos flhv35aflh, flhv35bflh, flhv33aflh, flhv33bflh, containing the gene targeting information, were added to pPRISM-V3-bactin RFP-DR48 using Golden Gate cloning as described in Wierson et al., 2018. The RFP cassette was liberated from the donor using the same *noto* sgRNA used to cut the genome. Gamma-crystalin:eGFP positive embryos were sorted and raised to adulthood, outcrossed to generate the F1 generation, and outcrossed again to generate lines of F2s.

### Southern blot analysis

Genomic Southern blot and copy number analysis was performed as described previously^42^. PCR primers used for genomic and donor specific probes are listed in Supplemental Table 2.

### Cell culture

HEK293 cells were obtained from ATCC (CRL-3216). Cells were maintained in Dulbecco’s Modified Eagle Medium (Gibco #11995-040) supplemented with 10% fetal bovine serum (Gibco #26140079) and 1% Penicillin Streptomycin (Gibco #15140-122). Media was changed every 2-3 days and replated at final dilution of 1:10 maintained at about 750,000 cells/ml.

### DNA isolation and Sequence analysis

DNA from whole cell populations was purified using Qiagen DNeasy Blood & Tissue Kit (Qiagen 69504). PCR amplification was performed with MyTaq DNA Polymerase (Bioline BIO-21108) and purified with Qiagen QIAquick PCR Purification kit (Qiagen 28104). Samples used for ICE analysis were submitted to GeneWiz Sanger Sequencing service.

### Cloning in vitro ErCas12a construct targeting AAVS1 (pErCas12aErCas12a-AAVS1)

Due to redundant restriction sites in the guide scaffold and ErCas12a protein, the pX601-AAV-CMV::NLS-SaCas9-NLS-3xHA-bGHpA;U6::BsaI-sgRNA plasmid (Addgene #61591) was digested with BsaI and NotI to first insert the ErCas12a secondary structure and sgRNA targeting AAVS1 with ErCas12a sgRNA AAVS1 top and ErCas12a sgRNA AAVS1 bottom (termed AAV: ErCas12a AAVS1 sgRNA) (Table 3). Following this ErCas12a as well as the Xenopus globin 5’ UTR and both N and C termini SV40 NLS signals were amplified from “T3TS nErCas12an” using PCR primers ErCas12a AgeI forward and ErCas12a BamHI reverse (Supplemental Table 1). The resulting 3.9kbp PCR fragment was cloned into an Agilent pSC Strataclone PCR cloning vector (termed ErCas12a Strataclone) to amplify the fragment with the desired restriction sites. ErCas12a Strataclone was digested with AgeI and BamHI to isolate ErCas12a with ends compatible with the AAV:ErCas12a AAVS1 sgRNA construct. The px601 plasmid was likewise digested with AgeI and BamHI to remove SaCas9 and replace it with ErCas12a. Plasmids were screened for insertion of both ErCas12a as well as the AAVS1 targeting sgRNA (pErCas12a-AAVS1) and amplified with Qiagen Endotoxin Free Maxiprep kit (Qiagen 12362).

### pErCas12a-AAVS1-U

pErCas12a-AAVS1-U was generated by inserting ErCas12a pre-U-crRNA into AAV 601 by digesting AAV601 with BsaI+ NotI and inserting annealed oligos ErCas12a UgRNA+Scaffold Top and ErCas12a UgRNA+Scaffold Bottom. The U6 promoter, the cr-RNA scaffold and the U-pre-crRNA were PCR amplified from AAV 601 using ErCas12a U6+ugRNA Primer PfoI Fw and ErCas12a U6+ugRNA Primer PfoI Rev. This PCR amplicon as well as pErCas12a-AAVS1 were digested with PfoI and ligated with T4 DNA ligase to generate pErCas12a-AAVS1-pre-crRNA1-U-pre-crRNA.

### Generating knock-in cassette for *AAVS1*

The 24 and 48bp homology arm CMV/GFP/Zeocin resistance knock-in cassettes were generated by designing PCR primers complementary to the psiRNA-SV40 Early PolyA GFPzeo plasmid (Invivogen) flanked by the 48 base pairs of homology and the UgRNA sequence separated by a 3bp spacer. The left 48HA forward primer AAVSI T1 L48HA and the right 48HA reverse primer AAVS1 T1 R48HA ^43^ were used to amplify the CMV/GFP/Zeocin resistance cassette containing the homology arms and the UgRNA target sequence (Supplemental Table 2, Supplemental Table 3). This 2.6kbp PCR fragment was subsequently cloned into a Strataclone PCR cloning vector and screened for the insert. Sequence confirmed knock in constructs were amplified with Qiagen Maxiprep Kit and termed “pGFP::Zeo-48”.

### Generating ErCas12a EZ Clone

nErCas12an was PCR amplified from pT3TSnErCas12an with Platinium Taq DNA polymerase HiFi (Thermo Fisher #11304011) with ErCas12a AgeI Fw and ErCas12a BamHI Rev and PCR cloned into the Agilent Strataclone vector (Agilent #240205) to generate ErCas12a Strataclone. ErCas12a Strataclone was subjected to site directed mutagenesis with ErCas12a SDM remove BsaI Top and ErCas12a SDM remove BsaI bottom with the Q5 SDM kit (New England Biolabs #E0554S) to generate ErCas12a Strataclone no BsaI. The nErCas12an fragment lacking the BsaI site was removed from the ErCas12a Strataclone no BsaI backbone by digesting with AgeI and BamHI. Likewise, the pX601-AAV-CMV::NLS-SaCas9-NLS-3xHA-bGHpA;U6::BsaI-sgRNA plasmid (Addgene #61591) was digested with AgeI and BamHI and had nErCas12an inserted into it to generate p601nErCas12an no BsaI. The cr-RNA scaffold was inserted into the p601nErCas12an no BsaI by digesting with BsaI and NotI and inserted the annealed oligos ErCas12a EZ clone scaffold top and ErCas12a EZ clone scaffold bottom to generate ErCas12a EZ Clone.

### Cloning in vitro ErCas12a construct targeting *CCR5* and *TRAC*

Cr-RNA oligos corresponding to the genomic target site (e.g. *CCR5* sgRNA 1 Top+ *CCR5* sgRNA 1 Bottom) were annealed in a thermocycler according to Zhang lab protocol. Briefly, oligos are annealed by incubating at 37°C for 30 minutes followed by 95°C for 5 minutes and then ramped down to 25°C at 5°C/min. The annealed oligo duplex was cloned into ErCas12a EZ clone Digested with BsaI and ligated by T4 DNA ligase.

### pMiniCAAGs:RFP-DR48 RFP assay generation, transfection, and analysis

pPRISM-V3-bactin SSA-DR48 was PCR amplified with Platinum Taq DNA polymerase HiFi (Thermo Fisher #11304011) with Broken RFP Transfer to Tol2 XhoI Fw and Broken RFP Transfer to Tol2 BglII Rev to add XhoI and BglII restriction sites to the RFP cassette. pkTol2C-EGFP as well as the RFP amplicon were digested with XhoI and BglII to create compatible cohesive ends and ligated with T4 DNA ligase to generate pkTol2CBrokenRFP. In order to for the construct to express episomally in cell culture systems a Kozak sequence was added by digesting pkTol2CBrokenRFP with EcoRI and XhoI and restriction cloning the annealed oligos Kozak Seq Top and Kozak Seq Bottom to generate pMiniCAAGs:RFP-DR48.

ErCas12a constructs targeting the UgRNA were generated by digesting ErCas12a EZ clone with BsaI and cloning in the annealed oligos ErCas12a UgRNA BsaI Top and ErCas12a UgRNA BsaI bottom to generate pErCas12a-U-pre-crRNA. Cas9 constructs targeting the UgRNA were generated by digesting lentiCRISPR v2 (Addgene #52961) with BsmBI and cloning in the annealed oligos Cas9 UgRNA BsmBI Top and Cas9 UgRNA BsmBI Bottom to generate pCas9-UgRNA. To control for promoter expression between ErCas12a and Cas9, the CMV promoter was added to pCas9-UgRNA by digesting ErCas12a EZ clone with XbaI and AgeI to remove the CMV promoter and pCas9-UgRNA with NheI and AgeI to replace the native EF1 alpha promoter.

HEK 293T cells were transfected with 5ug pMiniCAAGs:RFP-DR48 and 5ug of either pErCas12a-U-pre-crRNA or pCas9-UgRNA with the Etta H1 electroporator as described above. Cells were assessed for RFP expression 48 hours later by flow cytometry with 584nm emission and 607nm detection.

### Transfection

HEK293 cells used for indel acquisition assays were transfected with Liopofectamine 3000 (Thermo Fisher #L3000008) according to manufacturer’s protocol with 5ug of ErCas12a plasmid targeting each site. Cells used for targeted integration assays were transfected with Etta H1 electroporator with the following parameters: 200V, 784ms interval, 6 pulses, 1000us pulse duration, at a concentration of 20E6 cells/ml at the volume of 100ul in Etta EB electroporation buffer. Cells are recovered post electroporation by incubating at 37ºC for 5-10 minutes before being plated in a 6-well tissue culture plate at a density of about 1.5E6cells/ml.

### GeneWiz AmpliconEZ

#### DNA Library Preparation and Illumina Sequencing

DNA library preparations, sequencing reactions, and initial bioinformatics analysis were conducted at GENEWIZ, Inc. (South Plainfield, NJ, USA). DNA Library Preparation, clustering, and sequencing reagents were used throughout the process using NEBNext Ultra DNA Library Prep kit following the manufacturer’s recommendations (Illumina, San Diego, CA, USA). End repaired adapters were ligated after adenylation of the 3’ends followed by enrichment by limited cycle PCR. DNA libraries were validated on the Agilent TapeStation (Agilent Technologies, Palo Alto, CA, USA), and were quantified using Qubit 2.0 Fluorometer (Invitrogen, Carlsbad, CA) and multiplexed in equal molar mass. The pooled DNA libraries were loaded on the Illumina instrument according to manufacturer’s instructions. The samples were sequenced using a 2x 250 paired-end (PE) configuration. Image analysis and base calling were conducted by the Illumina Control Software on the Illumina instrument.

#### Data analysis

The raw Illumina reads were checked for adapters and quality via FastQC. The raw Illumina sequence reads were trimmed of their adapters and nucleotides with poor quality using Trimmomatic v. 0.36. Paired sequence reads were then merged to form a single sequence if the forward and reverse reads were able to overlap. The merged reads were aligned to the reference sequence and variant detection was performed using GENEWIZ proprietary Amplicon-EZ program.

#### Phylogeny and homology analysis

The amino acid sequence (ErCas12a) was used for homology search by Blastp tool (https://blast.ncbi.nlm.nih.gov/Blast.cgi) against the non-redundant (nr) database. The homologous sequences were then retrieved from NCBI database. Multiple sequence alignment was done using CLUSTAL Omega webserver (http://www.ebi.ac.uk/Tools/msa/clustalo/) The phylogenetic tree was constructed by using Maximum likelihood method based on JTT matrix-based model in MEGA 7.0.18 (Kumar, Stecher, and Tamura, 2016). Evaluation of branching was ensured by bootstrap statistical analysis (1000 replications).

#### Phylogenetic analysis

Phylogenetic tree relationship of ErCas12a from *Eubacterium rectale* and other related Cas proteins available in NCBI database was constructed by maximum likelihood method using MEGA7. The numbers above and below the branch points specify the confidence levels meant for the relationship of the paired sequences as estimated by the bootstrap analysis. Branch lengths are measured as the number of substitutions per site. Cas protein sequences used for phylogenetic analysis are CRISPR-associated endonuclease Cas12b from *Alicyclobacillus acidoterrestris*, CRISPR-associated Endonuclease Cas9 from *Streptococcus pyogens*, CRISPR-associated Endonuclease Cpf1 from *Acidaminococcus* sp. BV3L6, type V CRISPR-associated protein Cpf1 from *Lachnospiraceae* bacterium ND2006 and type V CRISPR-associated protein Cpf1 from *Francisella tularensis*.

#### Cas variant homology alignments

Multiple sequence alignment of ErCa12a with homologous Cas proteins like CRISPR-associated endonuclease Cas12b from *Alicyclobacillus acidoterrestris*, CRISPR-associated Endonuclease Cas9 from *Streptococcus pyogens*, Crispr-associated Endonuclease Cpf1 from *Acidaminococcus* sp. BV3L6, type V CRISPR-associated protein Cpf1 from *Lachnospiraceae* bacterium ND2006 and type V CRISPR-associated protein Cpf1 from *Francisella tularensis.* Six representative sequences were used for alignments with Clustal Omega webserver.

---

**Supplemental Figure 1.**
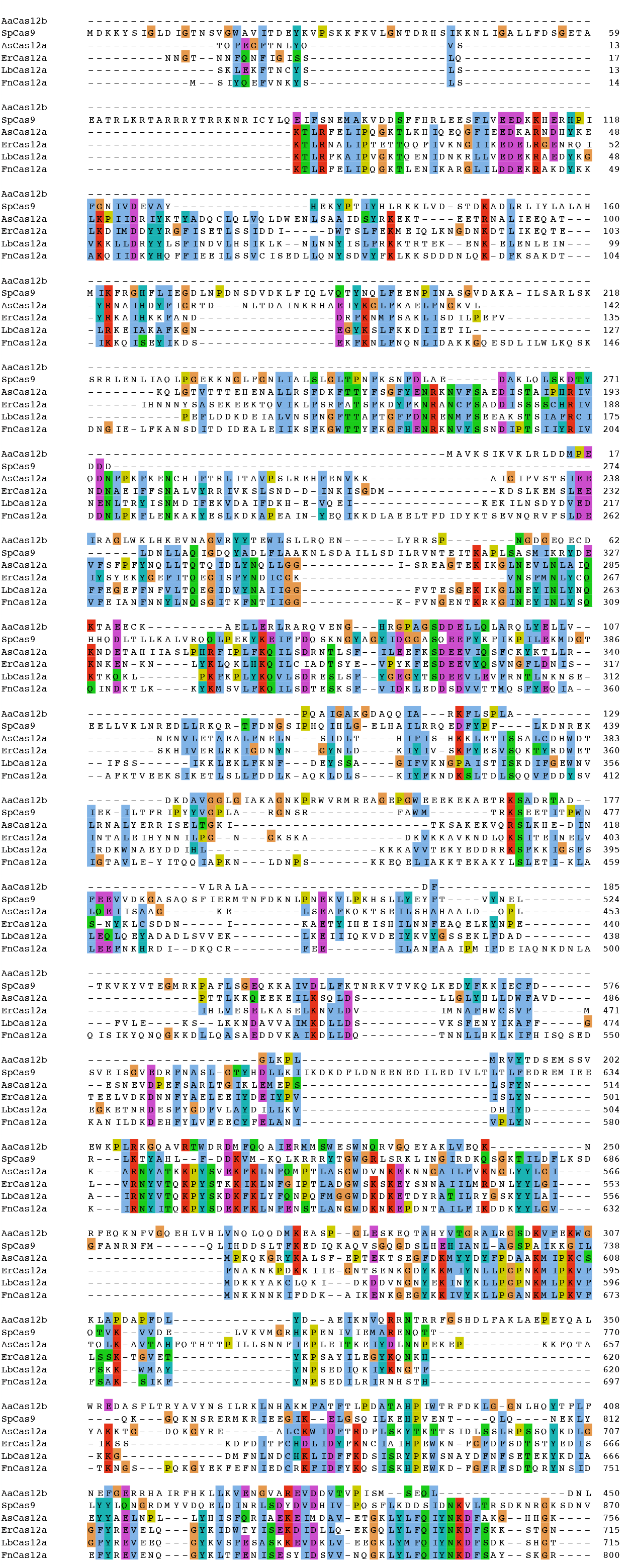
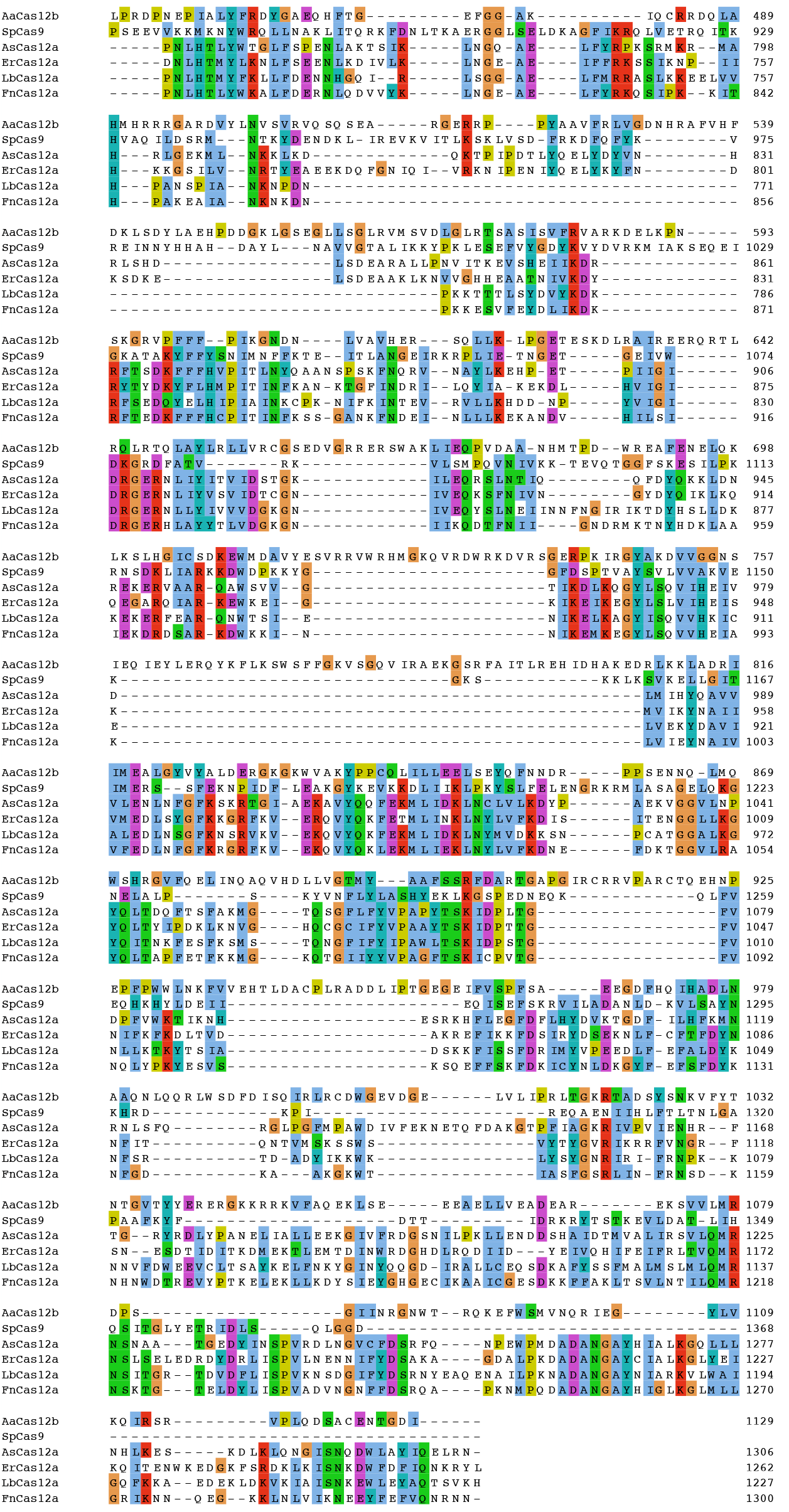
Amino acid alignment of Cas12a family members with SpCas9. Multiple sequence alignment of ErCa12a with homologous Cas proteins including CRISPR-associated endonuclease Cas12b from *Alicyclobacillus acidoterrestris*, CRISPR-associated Endonuclease Cas9 from *Streptococcus pyogens*, Crispr-associated Endonuclease Cpf1 from *Acidaminococcus* sp. BV3L6, type V CRISPR-associated protein Cpf1 from *Lachnospiraceae* bacterium ND2006 and type V CRISPR-associated protein Cpf1 from *Francisella tularensis.* Six representative sequences were used for alignments with Clustal Omega webserver.

**Supplemental Figure 2.**
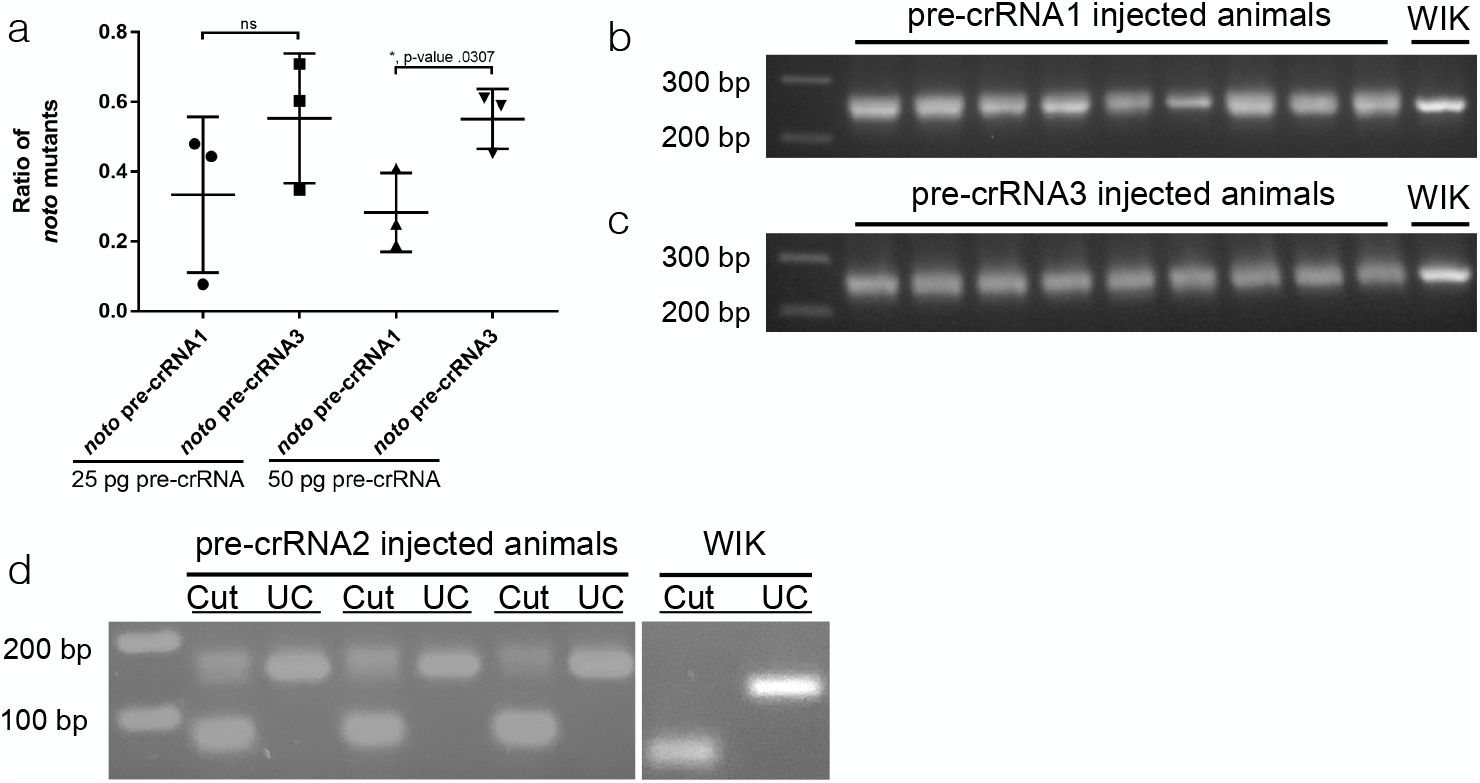
Qualitative representation of ErCas12a activity in zebrafish embryos. (a) Data plots showing ratio of injected animals displaying somatic *noto* phenotype. Data plots represent mean +/− s.d. p values calculated with one-tailed Student’s t-test. (b, c) Gel showing heteroduplex mobility shift of injected animals vs wild type (WIK) for pre-crRNA1 and pre-crRNA3 at *noto.* (d) RFLP analysis showing pre-crRNA2 is active at *cx43.4*.

**Supplemental Figure 3.**
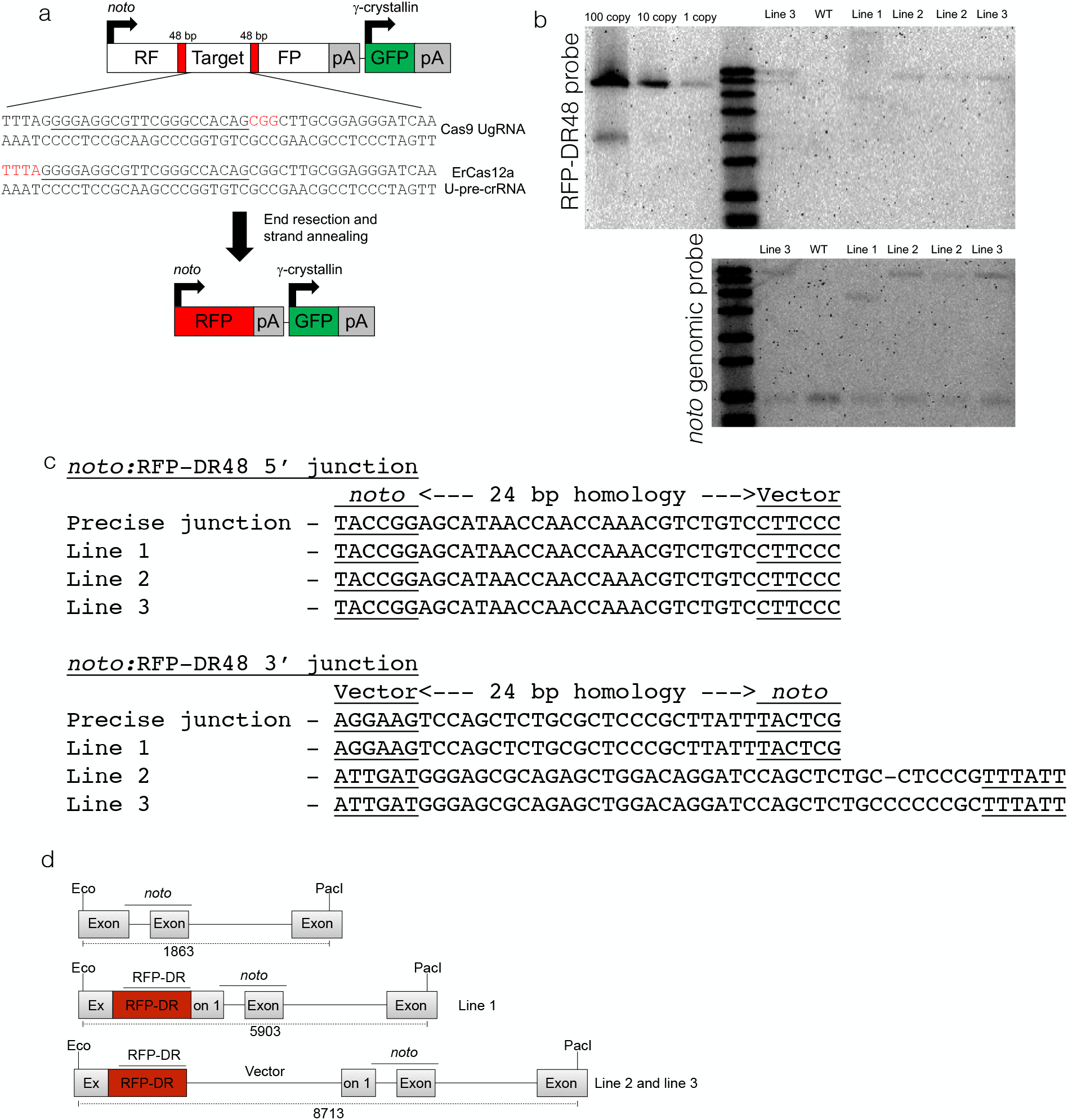
Engineering *noto*:RFP-DR48. (a) Schematic showing RFP-DR48 integrated at *noto*. The NBM (yellow) contains both SpCas9 and ErCas12a PAMs for universal targeting using the underlined RNA cursor target site. Red is the engineered stop codon. (b) Southern blot of three *noto*:RFP-DR48 lines. Band intensity is roughly single copy based on the copy number controls in the *noto* probed blot (top). Gel shifts are present indicating precise (line 1) and linear vector integration (lines 2 and 3). (c) Junction fragment analysis of integrations showing precision at the 5’ junction. Line 1 contains a precise 3’ integration, while line 2 and line 3 contain differing NHEJ events. (d) Schematic of integrations events as determined by Southern blot and DNA sequencing analysis.

**Supplemental Figure 4.**
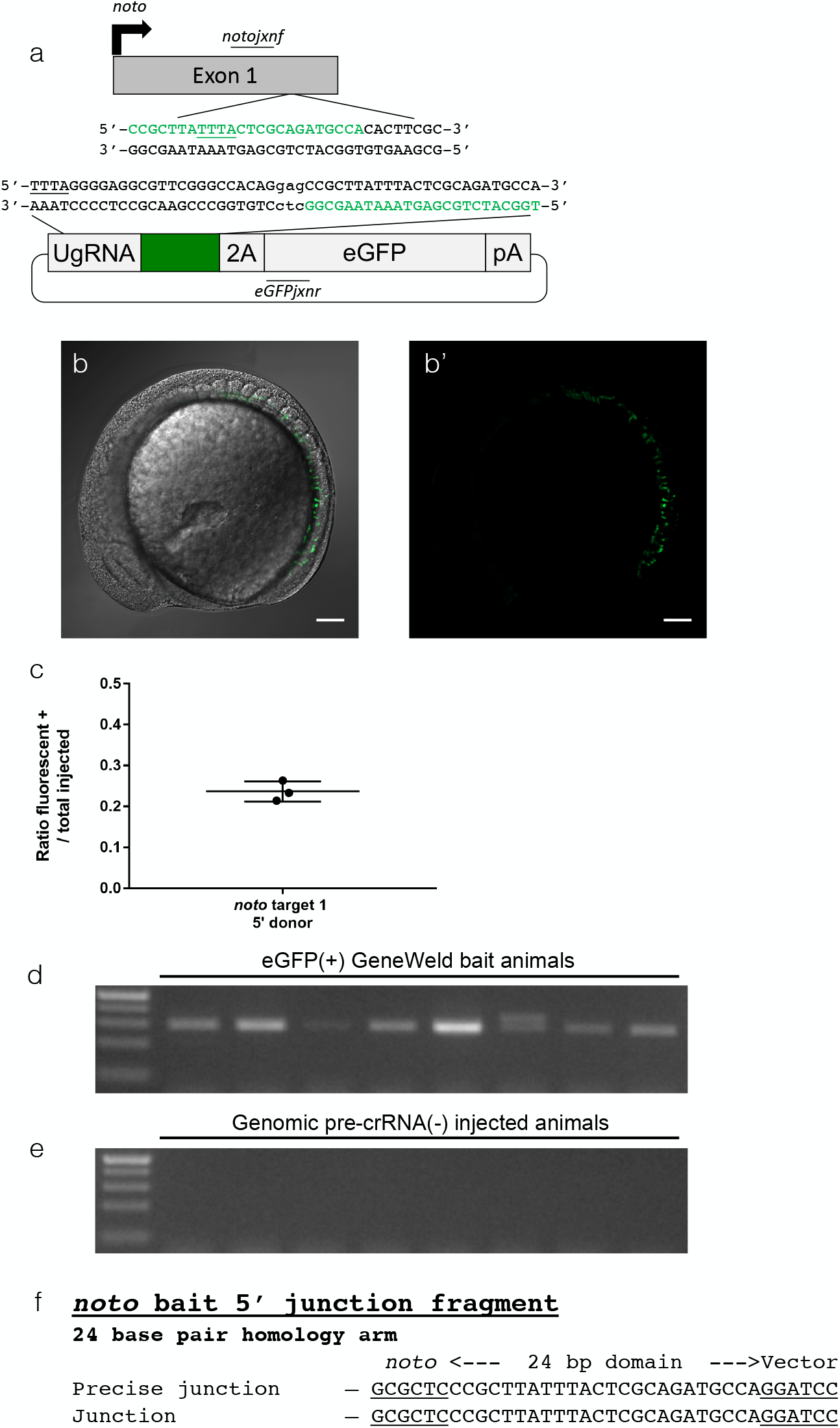
Targeting *noto* with a single homology domain. (a) Schematic showing designed homology for precise 5’ integration using ErCas12a. Green is designed homology. The PAM for ErCas12a targeting in the genome and donor is underlined. (b, b’) Representative confocal Z-stack image showing mosaic GFP expression in the notochord of an injected animal. Scale bar is 100 um. (c) Data plot showing the ratio of embryos with GFP expression in the notochord out of total injected embryos. Data plot represents mean +/− s.d. (d) Gel showing junction fragment expected after precise integration using the homology domain. (e) Gel showing no junction, indicating there is no integration without the genomic pre-crRNA. (f) DNA sequencing 5’ junctions showing precise integration using the programmed homology.

**Supplemental Figure 5.**
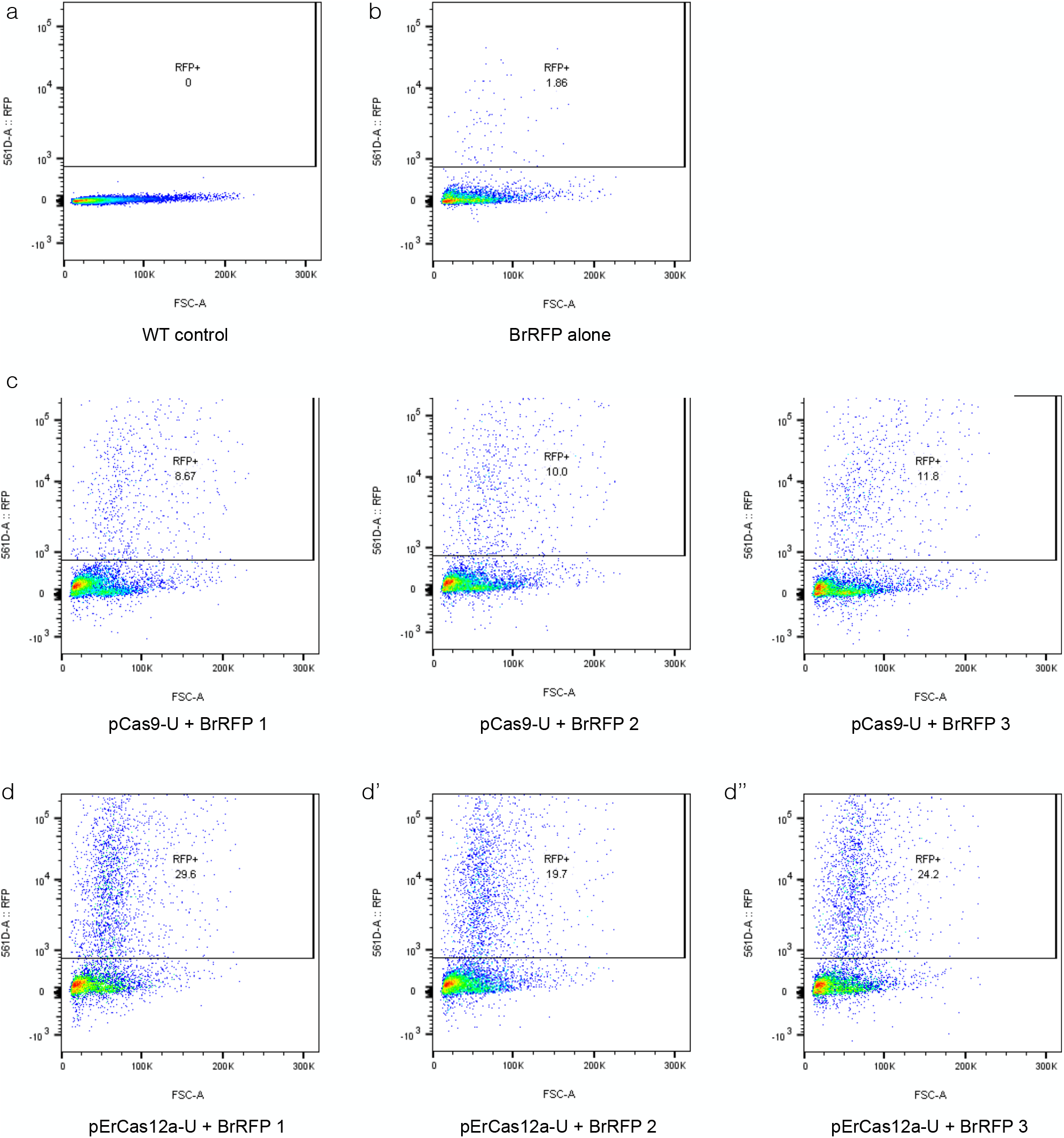
Raw data for flow cytometry of RFP-DR48 in HEK293 cells. (a) Untransfected HEK 293 cells (b) HEK 293 cells transfected with pMini-CAAGs::RFP-DR48 (c) HEK 293 cells transfected with SpCas9-UgRNA and pMini-CAAGs::RFP-DR48 (d) HEK 293 cells transfected with ErCas12a-pre-U-crRNA. The RFP plots shown were gated on the singlet cell population. All measurements taken using excitation of 584nm and emission of 607nm.

**Supplemental Figure 6.**
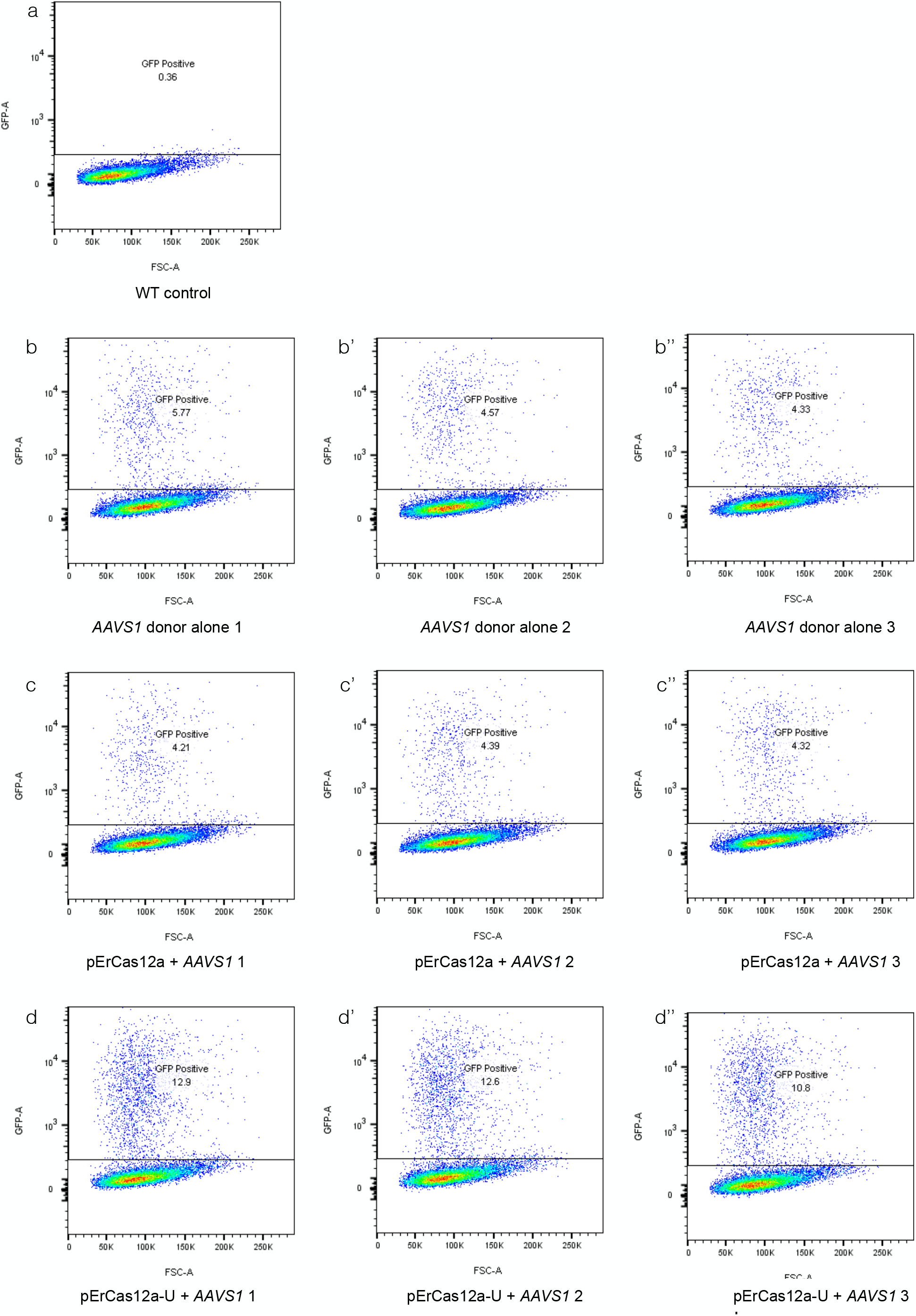
Raw data for flow cytometry of GeneWeld in HEK293 cells. (a) Untransfected HEK 293 cells (b-b’’) HEK 293 cells transfected only with the GFP GeneWeld donor (c-c’’) HEK 293 cells transfected with the GFP GeneWeld donor and pErCas12a-AAVS1 (d-d’’) HEK 293 cells transfected with the GFP GeneWeld donor and P-ErCas12a-U-AAVS1. The GFP plots shown were gated on the singlet cell population. All measurements taken using the FITC channel and excitation of 488nm and emission of 510nm.

