## Supplemental Tables for "Expanding the CRISPR Toolbox with ErCas12a in Zebrafish and Human Cells"

**Supplemental Table 1 - Mad7 cloning**

| gBlock name | Sequence |
|---|---|
| Fragment 1 | CCATGGCTTCTCCACCTAAGAAGAAGAGAAAGGTGAACAACGGAACTAATAATTTTC<br>AAAACTTCATTGGGATTAGTTCTCTGCAGAAGACCCTTCGGAATGCCCTCATTCCCA<br>CTGAGACGACTCAACAGTTTATCGTAAAAAATGGAATTATTAAGGAGGATG<br>AGTTGCGGGGGGAGAATAGGCAGATTTTGAAAGACATCATGGATGACTATTACCGGG<br>GTTTTATCTCCGAGACCCTGTCCTCTATCGATGATATTGATTGGACGTCTC<br>TTTTTGAGAAGATGGAGATTCAGCTGAAAAATGGTGATAACAAAGATACCCTCATTA<br>AGGAGCAAACCGAGTACCGGAAGGCGATCCACAAGAAGTTCGCCAACG<br>ATGATCGTTTTAAGAATATGTTCTCAGCCAAACTCATCAGTGACATCCTTCCAGAAT<br>TTGTAATTCATAATAATAACTACTCTGCGTCTGAGAAAGAAGAAAAAACTCA<br>AGTCATCAAGCTCTTTTCACGGTTTGCAACGAGCTTTAAGGATTACTTTAAAAACCG<br>CGCTAATTGTTTTTCTGCGGACGACATCAGCTCATCCAGCTGCCACAGAATC<br>GTCAATGACAATGCGGAGATCTTCTTCTCCAATGCTCTGGTATATAGGCGCATTGTA<br>AAGTCCTTGTCCAATGACGATATTAATAAGATAAGTGGTGATATGAAGGA<br>TTCTCTCAAGGAAATGTCATTGGAGGAGATCTACAGCTATGAGAAATACGGTGAATT<br>TATTACACAAGAAGGAATATCCTTTTATAATGACATCTGTGGGAAGGTGA ATTC |
| Fragment 2 | GAATTCTTTCATGAATTTGTACTGTCAAAAAAACAAGGAGAACAAAAACCTCTACAA<br>ATTGCAAAAACTGCATAAGCAAATTCTTTGTATAGCGGACACTAGCTATGATCC<br>TCGACAACATTTCCAGTAAGCATATCGTGGAACGGCTCAGGAAGATAGGG<br>GATAACTATAATGGCTATAACCTTGACAAGATCTATATCGTGAGTAAATTCTATGAA<br>AGTGTATCTCAAAAGACCTATCGAGATTGGGAAACCATAAACACAGCTCTT<br>GAGATTCATTACAATAATATTCTTCCTGGTAACGGGAAAAGTAAAGCCGATAAAGTG<br>AAGAAGGCCGTCAAAAACGACCTGCAGAAGAGCATAACGGAAATCAATG<br>AATTGGTGTCTAACTACAAGCTGTGCTCAGATGACAACATAAAAGCTGAGACATATA<br>TCCATGAGATCAGCCACATACTGAATAACTTTGAGGCGCAAGAGCTGAAAT<br>ATAATCCTGAGATCCACCTTGTAGAGTCTGAACTCAAGGCTTCCGAACTGAAAAATG<br>TACTTGACGTAATCATGAATGCTTTTCACTGGTGTAGTGTATTCATGACTGA<br>GGAACTGGTTGATAAGGATAATAATTTTTATGCGGAACTTGAAGAAATATACGATGA<br>GATTTATCCCGTTATCTCACTCTATAATTTGGTCCGAAATTATGTAACTCA<br>AAAACCATACTCCACAAAGAAAATCAAGCTCAATTTCGGTATCCCGACCTTGGCTGA<br>CGGATGGTCTAAAAGCAAGGAGTACTCCAATAACGCGATAATCTTGATGC<br>GAGATAATCTTTACTATCTCGGAATTTTTAATGCTAAAAATAAGCCCGATAAAAAGA<br>TTATTGAAGGAAACACATCTGAGAACAAAGGCGATTATAAAAAGATGATT<br>TATAATTTGCTCCCTGGACCAAACAAAATGATCCCTAAAGTTTTCCTCAGTTCCAAG<br>ACCGGGGTTGAGACGTACAAGCCTAGTGCATATATCTTGGAAGGTTATAAG<br>CAAAACAAGCACATCAAAAGTTCTAAGGACTTTGACATCACTTTTTGTCATGATTTG<br>ATTGACTATTTTAAAAACTGTATTGCAATTCACCCAGAGTGGAAGAATTTTG<br>GATTTGACTTCTCAGACACGTCTACCTATGAAGATATATCAGGATTTTATCGCGAGG<br>TTGAGCTCCAGGGTTACAAGATTGATTGGACTTATATCAGCGAGAAGGATA<br>TTGATCTTTTGCAGGAAAAAGGCCAACTTTATTTGTTCCAAATCTACAACAAGGACT<br>TTTCTAAGAAATCAACTGGCAACGATAACCTTCATACTATGTACCTCAAAAA<br>TCTCTTTTCCGAAGAGAATCTTAAGGATATCGTGCTCAAGCTGAACGGTGAGGCAGA<br>GATATTTTTCCGAAAGAGTTCTATCAAAAACCCAATTATCCACAAAAAAGG<br>CAGCATCCTGGTTAACAGGACGTACGGGTGCCGTATAAATTTGAGTCAGATGAGGAG<br>GTCTACCAAAGCGTAAACGGCT |

| | |
|---|---|
| Fragment 3 | CGTACGAGGCCGAAGAGAAAGATCAGTTCGGCAACATACAGATAGTGCGGAA<br>GAATATACCAGAGAATATCTACCAAGAGCTTTATAAGTATTTTAATGATAAGTCTGA<br>GCTACGGTTTTAAAAAAGGGAGGTTCAAGGTGGAGCGACAGGTGTACCA<br>AAAGTTTGAAACGATGCTTATTAATAAACTCAATTACCTCGTGTTCAAGGATATAAG<br>CATAACAGAAAATGGAGGGCTCCTTAAGGGATACCAGCTCACATACATAC<br>CGGACAAGCTTAAAAACGTGGGACACCAGTGCGGGTGTATATTTTACGTTCCTGCCG<br>CGTATACATCAAAGATAGACCCCACCACAGGGTTCGTGAATATCTTCAAG<br>TTTAAGGACTTGACAGTCGATGCAAAACGTGAGTTCATCAAGAAATTCGATTCAATC<br>CGGTACGATTCAGAAAAGAATCTGTTCTGTTTTACGTTCGATTATAACAACT<br>TTATTACGCAAAATACAGTGATGTCAAAGAGCTCATGGAGTGTCTACACATACGGGG<br>TTAGGATAAAGCGCAGGTTCGTTAACGGTCGGTTCTCAAACGAATCAGAC<br>ACGATTGACATTACGAAGGATATGGAAAAGACTCTGGAGATGACCGACATAAATTGG<br>CGAGACGGCCACGACCTCCGACAAGATATCATTGACTACGAGATCGTCC<br>AACACATTTTTGAAATCTTCCGGTTGACCGTCCAGATGCGAAACAGTCTTTCTGAAT<br>TGGAAGACCGGGATTACGACAGATTGATCAGTCCTGTATTGAACGAAAACA<br>ACATATTCTATGATTCCGCCAAAGCTGGCGATGCTTTGCCAAAAGACGCCGACGCGA<br>ATGGAGCATATTGTATCGCCCTTAAAGGCCTTTACGAAATCAAACAAATAA<br>CAGAGAACTGGAAAGAGGATGGGAAATTTAGCCGAGATAAGCTCAAGATCAGCAACA<br>AAGACTGGTTTGACTTTATTCAAAACAAACGGTACTTGCCGAAAAAGAA<br>ACGCAAAGTATAACCGCGGAGACAAAGAGCTTTCAGACGAGGCGGCGAAGTTGAAAA<br>ATGTAGTGGGACAT<br>CACGAAGCCGCCACAAACATCGTGAAGGACTATCGGTATACCTATGATAAGTACTTC<br>CTTCACATGCCAATCACGATCAATTTTAAAGCGAATAAGACCGGGTTCAT<br>AAATGACCGGATTCTGCAGTACATAGCAAAGGAGAAAGATCTTCATGTTATAGGCAT<br>TGATCGCGGCGAAAGAAACCTTATTTATGTCTCCGTTATAGACACATGTG<br>GGAACATCGTTGAACAAAAATCCTTTAATATCGTTAATGGATACGACTATCAGATAA<br>AGCTCAAACAACAGGAGGGGGCGCGCCAGATTGCTCGTAAAGAATGGAA<br>GGAAATAGGAAAAATAAAAGAAATCAAGGAGGGTTACCTGAGCCTTGTAATTCATGA<br>AATCTCCAAAATGGTTATAAAGTACAACGCGATTATTGTCATGGAAGATT |

| **Mad7 cloning primers** | **Sequence** |
|---|---|
| mad7f1 | ATGCCCATGGCTTCTCCACC |
| mad7r1 | GCATGAATTCACCTTCCCACAGATG |
| mad7f2 | ATGCGAATTCTTTCATGAATTTGTACTGTC |
| mad7r2 | GCATCGTACGTCCTGTTAACCAG |
| mad7f3 | ATGCCGTACGAGGCCGAAG |
| mad7r3 | GCATCCGCGGTTATACTTTGCG |

| **Mad7 cloning for human expression** | **Sequence** |
|---|---|
| Mad7 AgeI Fw | ACCGGTTTCTTTTTGCAGAAGCTCAG |
| Mad7 BamHIRev | GGATCCTACTTTGCGTTTCTTTTTCGG |
| Mad7 SDM remove BsaI Top | TTATCTCCGAGACGCTGTCCTCTAT |
| Mad7 SDM Remove BsaI Bottom | AACCCCGGTAATAGTCATCC |
| Mad7 EZ clone scaffold Top | CACCGTCAAAAGACCTTTTTAATTTCTACTCTTGTAGATAGAGACCGGGGGTGGTCT |
| Mad7 EZ clone scaffold Bottom | GGCCAAAAAaGAGACCacccccggtctctATCTACAAGAGTAGAAATTAAAAAGGTC |

**Supplemental Table 2 - Primers and oligos**

| Primer name | Sequence | Purpose |
|---|---|---|
| notojxnf | ATAGACGCTCTGCTCGCGAG | Indel and junction fragment analysis |
| notojxnr | CTTGTGCGTACACAGCTCCAC | Indel and junction fragment analysis |
| cx43.4jxnf1 | CCTTCGTGGGCAAGATATGGCTC | Indel and junction fragment analysis |
| cx43.4jxnf1r | TCTCACAACCAGGTTGCTGGG | Indel and junction fragment analysis |
| cx43.3jxnf2f | ATGGAAGAGATCGTGCCTGAGAAA | Indel and junction fragment analysis |
| cx43.4jxnf2r | AATCCTCGACAGAAGCTGCAGG | Indel and junction fragment analysis |
| *noto* site 1 | TTTACTCGCAGATGCCACACTTCGC | Mad7 target site |
| *noto* site 2 | GCGTACAGCCAAAGCATCATGCAAA | Mad7 target site |
| *noto* site 3 | GCGTACCGGAGCATAACCAACCAAA | Mad7 target site |
| *cx43.4* site 1 | AACATGACGGCCGGCGGAGAATAAA | Mad7 target site |
| *cx43.4* site 2 | TTTGACTGTTGTGGGGGGAGAATCG | Mad7 target site |
| SSA f | CGACATCCTGGCTACCAGCTTC | Indel and junction fragment analysis |
| SSA R | TGTCGCTTCTGCCTTCCAGG | Indel and junction fragment analysis |
| gfp5'R | GCTGAACTTGTGGCCGTTTA | Indel and junction fragment analysis |
| GFP3'F | ACATGGTCCTGCTGGAGTTC | Indel and junction fragment analysis |
| cxm7gRNA2fEZ | ATGCATGCCCTTCGTGGGCAAGATATGGCTC | Indel and junction fragment analysis |
| cxm7gRNA2rEZ | ATGCATGCTCTCACAACCAGGTTGCTGGG | Indel and junction fragment analysis |
| mad7noto1fEZ | CGTACGTAATAGACGCTCTGCTCGCGAG | Indel and junction fragment analysis |
| mad7noto1rEZ | CGTACGTACTTGTGCGTACACAGCTCCAC | Indel and junction fragment analysis |
| mad7noto3fEZ | TCGATCGAATAGACGCTCTGCTCGCGAG | Indel and junction fragment analysis |
| mad7noto3rEZ | TCGATCGACTTGTGCGTACACAGCTCCAC | Indel and junction fragment analysis |
| *noto* site 1 pre- crRNA | GTCAAAAGACCTTTTTAATTTCTACTCTTGTAGATCTCGCAGATGCCACACTTCGC | Synthego custom RNA |
| *noto* site 2 pre- crRNA | GTCAAAAGACCTTTTTAATTTCTACTCTTGTAGATCATGATGCTTTGGCTGTACGC | Synthego custom RNA |
| *noto* site 3 pre- crRNA | GTCAAAAGACCTTTTTAATTTCTACTCTTGTAGATGTTGGTTATGCTCCGGTACGC | Synthego custom RNA |
| *cx43.4* site 1 pre-crRNA | GTCAAAAGACCTTTTTAATTTCTACTCTTGTAGATTTCTCCGCCGGCCGTCATGTT | Synthego custom RNA |
| *cx43.4* site 2 pre-crRNA | GTCAAAAGACCTTTTTAATTTCTACTCTTGTAGATACTGTTGTGGGGGGAGAATCG | Synthego custom RNA |
| *noto* site 1 crRNA | GGAATTTCTACTCTTGTAGATCTCGCAGATGCCACACTTCGC | Synthego custom RNA |
| UgRNA pre-crRNA | GTCAAAAGACCTTTTTAATTTCTACTCTTGTAGATGGGGAGGCGTTCGGGCCACAG | Synthego custom RNA |
| v3f | ATGCCACCGGCGTTCCTAGTTCTTTAAACTGTTACAAAGTGTTTGG | RFP assay cloning |
| v3r | ATGCACTAGTTTCAATTAAGTTTGTGCCCCAGTTTGC | RFP assay cloning |
| bactinf | ACTAGTACGGACTGTTACCACTTCACGC | RFP assay cloning |
| bactinr | ATGCCGCCGGTGTAATTTATTTAGCAGTAGATAGCTATATTGTGTGAAACGC | RFP assay cloning |
| DRr | CCGCAAGCCGCTGTGGCCCGAACGCCTCCCCTAAATCACCCCTCTGATCTTGACGTT | RFP assay cloning |
| DRf | AGGGATCAAGCCTCCAGGACGGCTGC | RFP assay cloning |
| NBMf | CGACATCCTGGCTACCAGCTTC | Southern blot primers |
| NBMr | TGTCGCTTCTGCCTTCCAGG | Southern blot primers |
| flhSBf | CAGATGCCACACTTCGCGT | Southern blot primers |
| flhSBr | CGATGTTATACTTGCTTCTTTTTAGTTTTGTACATAT | Southern blot primers |
| flhv35aflh | TGATGGGAGCGCAGAGCTGGAGACAGGAAAAGCATAACCAACCAAACGCCTGTC | GeneWeld oligos |
| flhv35bflh | GAAGGACAGGCGTTTGGTTGGTTATGCTTTTCCTGTCTCCAGCTCTGCGCTCCC | GeneWeld oligos |
| flhv33aflh | AAGTCCAGCTCTGCGCTCCCGCTTATTGGGCCTGTCTCCAGCTCTGCGCTCCC | GeneWeld oligos |
| flhv33bflh | AGCGGGAGCGCAGAGCTGGAGACAGGCCCAATAAGCGGGAGCGCAGAGCTGGA | GeneWeld oligos |
| *noto* site 1 5' arm A 24 | AATTCTTTAGGGGAGGCGTTCGGGCCACAGGAGCCGCTTATTTACTCGCAGATGCCAG | GeneWeld oligos |
| *noto* site 1 5' arm B 24 | GATCCTGGCATCTGCGAGTAAATAAGCGGCTCCTGTGGCCCGAACGCCTCCCCTAAAG | GeneWeld oligos |
| *noto* site 3 5' arm A 24 | AATTCTTTAGGGGAGGCGTTCGGGCCACAGTTTGCGGAGATGAGAGAACGAACAAACG | GeneWeld oligos |
| *noto* site 3 5' arm B 24 | GATCCGTTTGTTCGTTCTCTCATCTCCGCAAACTGTGGCCCGAACGCCTCCCCTAAAG | GeneWeld oligos |
| *noto* site 3 3' arm A 24 | CATGGGAGCATAACCAACCAAACGCCTGTAAACTGTGGCCCGAACGCCTCCCCTAAAC | GeneWeld oligos |
| *noto* site 3 3' arm B 24 | GGCCGTTTAGGGGAGGCGTTCGGGCCACAGTTTACAGGCGTTTGGTTGGTTATGCTCC | GeneWeld oligos |
| **Human** | | |
| *AAVS1* site *1* | TTTAGGACGGTGCGATCCCACCCCT | |
| *AAVS1* site 2 | TTTCCCTCCAGGACTTGTCCAAGGA | |
| *CCR5* site 1 | TTTACAGGAAACCCATAGAAGACAT | |
| *CCR5* site 2 | TTTCGCCTTCAATACACTTAATGAT | |
| *CCR5* site 3 | GGGCTTTTGACTAGATGAATGTAAA | |
| *CCR5* site 4 | TAGTTAGCTTCTGAGATGAGTAAAA | |
| *TRAC* site 1 | TTTTGATTCTCAAACAAATGTGTCA | |
| *TRAC* site 2 | TCTGATGTGTATATCACAGACAAAA | |
| *TRAC* site 3 | CAGTGCTGTGGCCTGGAGCAACAAA | |
| *TRAC* site 4 | TTTGCATGTGCAAACGCCTTCAACA | |
| *AAVSI* T1 L48HA | CTTTAGGACGGTGCGATCCCACCCCTCTGAGCCAGAATCGGAAGAGCCAGACGGAGGCTTTAGGACGGTGCGATCCTGCAGGCGTTACATAACTT | Homology arm oligo |
| *AAVS1* T1 R48HA | CTTTAGGACGGTGCGATCCCACCCCTGGGAAGCCAGGAAGCTGGGCACCCAGCAGGCAAAGCCCCGACGGAGGGTTACGGTTCCTGGCCTTTTGCTG | Homology arm oligo |
| *AAVS1* T1 L24HA | CTTTAGGACGGTGCGATCCCACCCCTCGGAGGCTTTAGGACGGTGCGATCCCTGCAGGCGTTACATAACTT | Homology arm oligo |
| *AAVS1* T1 R24HA | CTTTAGGACGGTGCGATCCCACCCCTACCCAGCAGGCAAAGCCCCGACGGTTACGGTTCCTGGCCTTTTGCTG | Homology arm oligo |
| *AAVS1* L48HA+UgRNA+3bp spacer | TTTAGGGGAGGCGTTCGGGCCACAGGTTTGAGCCAGAATCGGAAGAGCCAGACGGAGGCTTTAGGACGGTGCGATCCCTGCAGGCGTTACATAACT | Homology arm oligo |
| *AAVS1* R48HA+UgRNA+3bp Spacer | TTTAGGGGAGGCGTTCGGGCCACAGCTTGGCGGGGAAGCCAGGAAGCTGGGCACCCAGCAGGCAAAGCCCCGACGGTTACGGTTCCTGGCCTTTTGCT | Homology arm oligo |
| *AAVS1* Target1FW | CCCATTGAACCCCCGTCTAC | Indel and junction fragment analysis |
| *AAVS1* Target1Rev | TGCCCCTACTCACGAATCTC | Indel and junction fragment analysis |
| *AAVS1* ExonTargetFw | TGCAGCTTCGGAACCAAAAAG | Indel and junction fragment analysis |
| *AAVS1* ExonTargetRev | GGGGCAGTTCCCTCGACTG | Indel and junction fragment analysis |
| *TRAC* Fw | GAGCAGCTGGTTTCTAAGATGC | Indel and junction fragment analysis |
| *TRAC* Rev | GGAGAGGCAACTTGGAGAAGG | Indel and junction fragment analysis |
| *AAVS1* NGS Fw | ACACTCTTTCCCTACACGACGCTCTTCCGATCTAGTCCCTCCCATACCCATTGA | Indel and junction fragment analysis |
| *AAVS1* NGS Rev | GACTGGAGTTCAGACGTGTGCTCTTCCGATCTGGTGGAGGATGGGGCGATG | Indel and junction fragment analysis |
| *CCR5* Fw | GTTAATGTGAAGTCCAGGATCCC | Indel and junction fragment analysis |
| *CCR5* Rev | TCTGCAAATCTTTCTTTTGAGAGGT | Indel and junction fragment analysis |
| CMVUnivRev | CCCGTGAGTCAAACCGCTAT | Indel and junction fragment analysis |
| Mad7 sgRNA AAVS1 top | CACCGTCAAAAGACCTTTTTAATTTCTACTCTTGTAGATGGACGGTGCGATCCCACCCCTTTTTT | IDT Oligos |
| Mad7 sgRNA AAVS1 bottom | GGCCAAAAAAGGGGTGGGATCGCACCGTCCATCTACAAGAGTAGAAATTAAAAAGGTCTTTTGAC | IDT Oligos |
| *AAVS1* Exon Target Top Oligo | AGATCCTCCAGGACTTGTCCAAGGA | IDT Oligos |
| *AAVS1* Exon Target Bottom Oligo | AAAATCCTTGGACAAGTCCTGGAGG | IDT Oligos |
| *CCR5* sgRNA 1 Top | AGATCAGGAAACCCATAGAAGACAT | IDT Oligos |
| *CCR5* sgRNA 1 Bottom | AAAAATGTCTTCTATGGGTTTCCTG | IDT Oligos |
| *CCR5* sgRNA 2 Top | AGATGCCTTCAATACACTTAATGAT | IDT Oligos |
| *CCR5* sgRNA 2 Bottom | AAAAATCATTAAGTGTATTGAAGGC | IDT Oligos |
| *CCR5* sgRNA 3 Top | AGATCATTCATCTAGTCAAAAGCCC | IDT Oligos |
| *CCR5* sgRNA 3 Bottom | AAAAGGGCTTTTGACTAGATGAATG | IDT Oligos |
| *CCR5* sgRNA 4 Top | AGATACTCATCTCAGAAGCTAACTA | IDT Oligos |
| *CCR5* sgRNA 4 Bottom | AAAATAGTTAGCTTCTGAGATGAGT | IDT Oligos |
| *TRAC* sgRNA 1 Top | AGATGATTCTCAAACAAATGTGTCA | IDT Oligos |
| *TRAC* sgRNA 1 Bottom | AAAATGACACATTTGTTTGAGAATC | IDT Oligos |
| *TRAC* sgRNA 2 Top | AGATTCTGATGTGTATATCACAGAC | IDT Oligos |
| *TRAC* sgRNA 2 Bottom | AAAAGTCTGTGATATACACATCAGA | IDT Oligos |
| *TRAC* sgRNA 3 Top | AGATCAGTGCTGTGGCCTGGAGCAA | IDT Oligos |
| *TRAC* sgRNA 3 Bottom | AAAATTGCTCCAGGCCACAGCACTG | IDT Oligos |
| *TRAC* sgRNA 4 Top | AGATCATGTGCAAACGCCTTCAACA | IDT Oligos |
| *TRAC* sgRNA 4 Bottom | AAAATGTTGAAGGCGTTTGCACATG | IDT Oligos |
| Broken RFP Transfer to Tol2 XhoI Fw | GGTCCTCTCGAGGTGTCTAAG | RFP assay cloning |
| Broken RFP Transfer to Tol2 BglII Rev | AAAAGATCTTCAATTAAGTTTGTGCCCCAGT | RFP assay cloning |
| Kozak Seq Top | AATTAGCCACCATGG | RFP assay cloning |
| Kozak Seq Bottom | TCGACCATGGTGGCT | RFP assay cloning |
| Mad7 UgRNA BsaI Top | AGATGGGGAGGCGTTCGGGCCACAG | IDT Oligos |
| Mad7 UgRNA BsaI Bottom | AAAACTGTGGCCCGAACGCCTCCCC | IDT Oligos |
| Cas9 UgRNA BsmBI Top | CACCGGGAGGCGTTCGGGCCACAG | IDT Oligos |
| Cas9 UgRNA BsmBI Bottom | AAACCTGTGGCCCGAACGCCTCCC | IDT Oligos |
| Mad7 UgRNA+Scaffold Top | CACCGTCAAAAGACCTTTTTAATTTCTACTCTTGTAGATGGGGAGGCGTTCGGGCCACAGTTTTTggcc | IDT Oligos |
| Mad7 UgRNA+Scaffold Bottom | GGCCAAAAACTGTGGCCCGAACGCCTCCCCATCTACAAGAGTAGAAATTAAAAAGGTCTTTTGACggtg | IDT Oligos |
| Mad7 U6+ugRNA Primer Pfo Fw | AAATCCGGGAGAGGGCCTATTTCCCATGAT | IDT Oligos |
| Mad7 U6+ugRNA Primer Pfo1 Rev | AAATCCcGGAGCGGCCAAAAACTGTGGCCC | |

**Supplemental Table 3 - Targeting domain information for all gene targeting experiments. N/A means not applicable**

| Genomic target | Donor vector | Genomic target sequence | Donor target sequence | 5' spacer | 5' homology arm | 3' spacer | 3' homology arm |
|---|---|---|---|---|---|---|---|
| Zebrafish | Zebrafish | | | | | | |
| *noto* site 1 | p494-2a:eGFP-pA | TTTACTCGCAGATGCCACACTTCGC | TTTAGGGGAGGCGTTCGGGCCACAG | gag | CCGCTTATTTACTCGCAGATGCCA | N/A | N/A |
| *noto* site 3 | p494-2a:eGFP-pA | AAACGCGTACCGGAGCATAACCAAC | TTTAGGGGAGGCGTTCGGGCCACAG | ttt | GCGGAGATGAGAGAACGAACAAAC | aaa | GAGCATAACCAACCAAACGCCTGT |
| *noto* | pPRISM-V3:RFP-DR | GGGAGCGCAGAGCTGGAGACAGG | GGGAGCGCAGAGCTGGAGACAGG | aaa | AGCATAACCAACCAAACGCCTGTC | ggg | TCCAGCTCTGCGCTCCCGCTTATT |
| Human | Human cells | | | | | | |
| AAVS1 site 1 | pCMV:GFP::Zeo-48 | TTTAGGACGGTGCGATCCCACCCCT | TTTAGGGGAGGCGTTCGGGCCACAG | gtt | TGAGCCAGAATCGGAAGAGCCAGACGGAGGCTTTAGGACGGTGCGATC | aag | CCGTCGGGGCTTTGCCTGCTGGGTGCCCAGCTTCCTGGCTTCCCCGCC |

**Supplemental Table 4 - GeneWeld results**

| Genomic target | Donor vector | Donor sgRNA target (genomic or UgRNA) | Homology length (5'/3') | Experiment number | Reporter positive embryos | Total embryos | Percent with positive report | Standard Error |
|---|---|---|---|---|---|---|---|---|
| *noto* site 1 | p494-2a:eGFP-pA | UgRNA | 24/x | 1 | 21 | 80 | 26.30% | |
| | p494-2a:eGFP-pA | UgRNA | 24/x | 2 | 18 | 84 | 21.40% | |
| | p494-2a:eGFP-pA | UgRNA | 24/x | 3 | 17 | 73 | 23.30% | |
| | | | | Average | 56 | 237 | 23.60% | 0.01146295 |
| *noto* site 3 | p494-2a:eGFP-pA | UgRNA | 24/24 | 1 | 33 | 74 | 44.60% | |
| | p494-2a:eGFP-pA | UgRNA | 24/24 | 2 | 9 | 50 | 18.00% | |
| | p494-2a:eGFP-pA | UgRNA | 24/24 | 3 | 14 | 57 | 24.60% | |
| | | | | Average | 56 | 181 | 30.90% | 0.06530989 |

Supplemental Table 5 - RFP-DR48 results

| Injection Date | Line | Nuclease | UgRNA | 4 hour post injection incubation temperature | Injected animals | Injected animals gammy-cry + | RFP + | RFP+/Total | Broad | Intermediate | Narrow | Broad % of positive | Intermediate % of positive | Narrow % of positive |
|---|---|---|---|---|---|---|---|---|---|---|---|---|---|---|
| 6/6/18 | F1-Male1:F1-Female2:F2 outx | Mad7 300 pg mRNA | 25 pg synthego pre-crRNA | 34ºC | 39 | 18 | 13 | 0.722 | 2 | 6 | 5 | 0.153846154 | 0.461538462 | 0.384615385 |
| 6/13/18 | F1-Male1:F1-Female2:F2 outx | cas9 150 pg mRNA | 25 pg IDT | 28ºC | 34 | 18 | 4 | 0.222 | 0 | 2 | 2 | 0 | 0.5 | 0.5 |
| 6/19/18 | F1-Male1:F1-Female2:F2 outx | Cas9 300 pg mRNA | 25 pg IDT | 28ºC | 39 | 20 | 7 | 0.350 | 3 | 4 | 0 | 0.428571429 | 0.571428571 | 0 |
| | F1-Male1:F1-Female2:F2 outx | Cas9 300 pg mRNA | 25 pg IDT | 28ºC | 25 | 14 | 7 | 0.500 | 3 | 2 | 2 | 0.428571429 | 0.285714286 | 0.285714286 |
| | F1-Male1:F1-Female2:F2 outx | Mad7 300 pg mRNA | 25 pg synthego pre-crRNA | 34ºC | 63 | 32 | 28 | 0.875 | 4 | 10 | 14 | 0.142857143 | 0.357142857 | 0.5 |
| | F1-Male1:F1-Female2:F2 outx | Mad7 300 pg mRNA | 25 pg synthego pre-crRNA | 34ºC | 26 | 15 | 11 | 0.733 | 0 | 7 | 4 | 0 | 0.636363636 | 0.363636364 |
| 6/25/18 | F1-Male1:F1-Female2:F2 outx | Mad7 300 pg mRNA | 25 pg synthego pre-crRNA | 34ºC | 95 | 48 | 28 | 0.583 | 3 | 16 | 9 | 0.107142857 | 0.571428571 | 0.321428571 |
| 6/26/18 | F1-Male1:F1-Female2:F2 outx | Cas9 300 pg mRNA | 25 pg IDT | 28*C | 44 | 27 | 11 | 0.407 | 4 | 5 | 2 | 0.363636364 | 0.454545455 | 0.181818182 |
| 6/26/18 | F1-Male1:F1-Female2:F2 outx | Cas9 300 pg mRNA | 25 pg IDT | 34ºC | 57 | 27 | 9 | 0.333 | 4 | 4 | 1 | 0.444444444 | 0.444444444 | 0.111111111 |
| | F1-Male1:F1-Female2:F2 outx | Cas9 300 pg mRNA | 25 pg IDT | 28ºC | 67 | 32 | 12 | 0.375 | 5 | 3 | 4 | 0.416666667 | 0.25 | 0.333333333 |
| 8/13/18 | F1-Male1:F1-Female2:F2 outx | Cas9 300 pg mRNA | 25 pg IDT | 28ºC | 109 | 57 | 25 | 0.439 | 6 | 11 | 8 | 0.24 | 0.44 | 0.32 |
| | F1-Male1:F1-Female2:F2 outx | Mad7 300 pg mRNA | 25 pg synthego pre-crRNA | 34ºC | 119 | 53 | 44 | 0.830 | 9 | 21 | 14 | 0.20454545 | 0.47727273 | 0.31818182 |
| 10/3/18 | F1-Male1:F1-Female2:F2 outx | Mad7 300 pg mRNA | 25 pg synthego pre-crRNA | 34ºC | 104 | 49 | 29 | 0.592 | 7 | 14 | 8 | 0.24137931 | 0.48275862 | 0.27586207 |
| | F1-Male1:F1-Female2:F2 outx | Cas9 300 pg mRNA | 25 pg IDT | 28ºC | 110 | 50 | 16 | 0.320 | 4 | 8 | 4 | 0.25 | 0.5 | 0.25 |

Supplemental Table 6 - ICE analysis of HEK293

| Sample | ICE | KO-Score | ICE d | R Squared | Mean Discord Before | Mean Discord After | Seq Primer |
|---|---|---|---|---|---|---|---|
| CCR51_1 | 62 | 43 | 65 | 0.97 | 0.093 | 0.464 | TCTGCAAATCTTTCTTTTGAGAGGT |
| CCR51_2 | 46 | 30 | 48 | 0.98 | 0.139 | 0.358 | TCTGCAAATCTTTCTTTTGAGAGGT |
| CCR51_3 | 60 | 42 | 63 | 0.96 | 0.122 | 0.46 | TCTGCAAATCTTTCTTTTGAGAGGT |
| CCR5_2_1 | 51 | 27 | 50 | 0.96 | 0.047 | 0.458 | GCCTTACTGTTGAAAAGCCCTG |
| CCR5_2_2 | 50 | 28 | 48 | 0.97 | 0.041 | 0.327 | GCCTTACTGTTGAAAAGCCCTG |
| CCR5_2_3 | 50 | 29 | 48 | 0.97 | 0.036 | 0.333 | GCCTTACTGTTGAAAAGCCCTG |
| CCR5_3_1 | 31 | 25 | 28 | 0.97 | 0.069 | 0.21 | GCCTTACTGTTGAAAAGCCCTG |
| CCR5_3_2 | 31 | 24 | 30 | 0.97 | 0.082 | 0.235 | GCCTTACTGTTGAAAAGCCCTG |
| CCR5_3_3 | 27 | 22 | 24 | 0.98 | 0.06 | 0.203 | GCCTTACTGTTGAAAAGCCCTG |
| CCR54_1 | 24 | 19 | 23 | 0.99 | 0.097 | 0.282 | TCTGCAAATCTTTCTTTTGAGAGGT |
| CCR54_2 | 22 | 19 | 21 | 0.99 | 0.088 | 0.21 | TCTGCAAATCTTTCTTTTGAGAGGT |
| CCR54_3 | 22 | 18 | 20 | 0.99 | 0.082 | 0.206 | TCTGCAAATCTTTCTTTTGAGAGGT |
| TRAC_1_1 | 13 | 13 | 13 | 0.99 | 0.102 | 0.399 | GAGCAGCTGGTTTCTAAGATGC |
| TRAC_1_2 | 19 | 17 | 19 | 0.98 | 0.125 | 0.212 | GAGCAGCTGGTTTCTAAGATGC |
| TRAC_1_3 | 20 | 18 | 21 | 0.99 | 0.12 | 0.215 | GAGCAGCTGGTTTCTAAGATGC |
| TRAC_2_1 | 63 | 35 | 68 | 0.96 | 0.124 | 0.518 | GAGCAGCTGGTTTCTAAGATGC |
| TRAC_2_2 | 59 | 33 | 63 | 0.96 | 0.123 | 0.462 | GAGCAGCTGGTTTCTAAGATGC |
| TRAC_2_3 | 57 | 34 | 61 | 0.96 | 0.109 | 0.438 | GAGCAGCTGGTTTCTAAGATGC |
| TRAC_3_1 | 51 | 38 | 53 | 0.96 | 0.125 | 0.377 | GAGCAGCTGGTTTCTAAGATGC |
| TRAC_3_2 | 50 | 38 | 53 | 0.95 | 0.134 | 0.395 | GAGCAGCTGGTTTCTAAGATGC |
| TRAC_3_3 | 46 | 36 | 48 | 0.96 | 0.119 | 0.459 | GAGCAGCTGGTTTCTAAGATGC |
| TRAC_4_1 | 32 | 22 | 34 | 0.97 | 0.137 | 0.244 | GAGCAGCTGGTTTCTAAGATGC |
| TRAC_4_2 | 33 | 22 | 35 | 0.96 | 0.146 | 0.262 | GAGCAGCTGGTTTCTAAGATGC |
| TRAC_4_3 | 29 | 20 | 29 | 0.97 | 0.113 | 0.225 | GAGCAGCTGGTTTCTAAGATGC |
| AAVS1_1 | 54 | 45 | 47 | 0.98 | 0.057 | 0.42 | CCCATTGAACCCCCGTCTAC |
| AAVS1_2 | 54 | 46 | 47 | 0.98 | 0.052 | 0.448 | CCCATTGAACCCCCGTCTAC |
| AAVS1_3 | 66 | 56 | 59 | 0.98 | 0.06 | 0.486 | CCCATTGAACCCCCGTCTAC |
